# High-Dimensional Mediation Analysis with Network Mediators: Applications to Pediatric Acute Lymphoblastic Leukemia

**DOI:** 10.1101/2024.09.23.614601

**Authors:** Jade Xiaoqing Wang, Zhao-Hua Lu, Wilburn E Reddick, Heather M Conklin, John O Glass, Lisa Jacola, Arzu Onar-Thomas, Sima Jeha, Cheng Cheng, Xiang Zhou, Yimei Li

**Author notes:** Correspondence to XZ and YL.

## Abstract

Acute lymphoblastic leukemia (ALL) is the most common childhood cancer, with survivors frequently experiencing long-term neurocognitive morbidities. Here, we utilize the TOTXVI clinical trial data to elucidate the mechanisms underlying treatment-related neurocognitive side effects in pediatric ALL patients by incorporating brain connectivity network data. To enable such analysis, we propose a high-dimensional mediation analysis method with a novel network mediation structural shrinkage (NMSS) prior, which is particularly suited for analyzing high-dimensional brain structural connectivity network data that serve as mediators. Our method is capable of addressing the structural dependencies of brain connectivity networks including sparsity, effective degrees of nodes, and modularity, yielding accurate estimates of the high-dimensional coefficients and mediation effects. We demonstrate the effectiveness and superiority of the proposed NMSS method through simulation studies and apply it to the TOTXVI data, revealing significant mediation effects of brain connectivity on visual processing speed directed by IT intensity. The findings shed light on the potential of targeted interventions to mitigate neurocognitive deficits in pediatric ALL survivors.

## 1 INTRODUCTION

Acute lymphoblastic leukemia (ALL) is the most common childhood cancer, accounting for 25% of all cancers in children and roughly 75-80% of all pediatric leukemia cases (Pui, Nichols and Yang, 2019). The success in treating childhood ALL has resulted in a growing number of survivors, with the 5-year overall survival rate exceeding 90%. However, such success also highlights the needs to investigate and address the long-term neurocognitive morbidities for the large number of survivors. Specifically, current treatment protocols for ALL employ systemic chemotherapy in conjunction with intrathecal (IT) chemotherapy for central nervous system (CNS)-directed prophylaxis, effectively sparing most patients from the significant and enduring neurocognitive declines associated with cranial irradiation (Harila et al., 2009). Nevertheless, pediatric patients remain at risk for neurocognitive deficits due to their vulnerability particularly in areas such as processing speed, attention, and executive functioning (Iyer et al., 2015). Therefore, it becomes critically important to carry out prospective integrative analysis to elucidate the mechanisms underlying these side effects, quantify their impact, and identify causal risk factors, ultimately improving treatment protocols and the quality of life for ALL survivors.

A particular motivating example that enables such integrative analysis is the TOTXVI clinical trial (ClinicalTrials.gov identifiers: NCT00549848), a phase III randomized treatment protocol based on intensified systemic chemotherapy and optimized IT chemotherapy for pediatric ALL. Comprehensive patient data were collected during the study, including treatment-related clinical information, brain Diffusion Tensor Imaging (DTI) data, and neurocognitive assessments. The brain DTI data were preprocessed into undirected structural connectivity networks (Sotiropoulos and Zalesky, 2019) depicting the interconnections of brain regions through fiber bundles. Previous studies in other trials have shown that IT chemotherapy can lead to structural brain abnormalities such as white matter damage and leukoencephalopathy (Cheung et al., 2016), while structural brain abnormalities have been linked to neurocognitive impairments in ALL survivors (Cheung and Krull, 2015). Several methods have also been developed to perform pairwise associations using high-dimensional brain connectivity either as predictors (Guha and Guhaniyogi, 2021; Zhang et al., 2019) or outcomes (Guha and Guhaniyogi, 2021; Roy et al., 2019). However, few studies have explored the full causal pathway from treatment variables through brain connectivity to neurocognitive outcomes, which is key for understanding the brain structural mechanism underlying IT chemotherapy effects on neurocognitive impairments.

To better understand the causal mechanisms of treatment side effects on neurocognitive outcomes, we obtained n=271 TOTXVI participants with complete treatment, DTI, and neurocognitive data. Such data presents a unique opportunity for us to integrate brain structural connectivity measurements through mediation analysis to elucidate the potentially causal pathway from IT chemotherapy (exposure variable) to brain connectivity developments (mediators) and subsequently to neurocognitive deficits (outcome variable; depicted by the pathway in red in Figure 1a).

**Fig 1:**
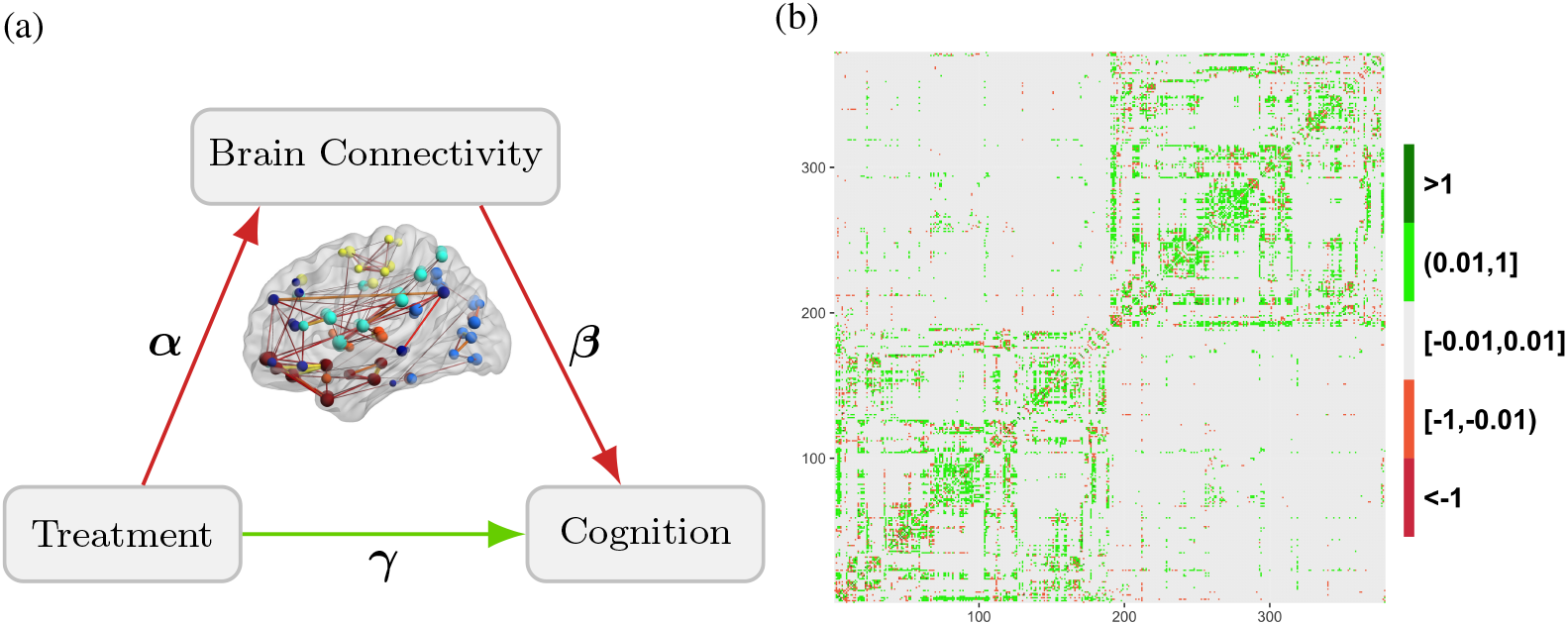
(a) Path diagram of the mediation analysis model with treatment (exposure), brain connectivity (mediator), and cognition (outcome). (b) A processed connectivity matrix in TOTXVI real data analysis in Section 6.

Mediation analysis (MacKinnon, 2008) is a general framework for revealing causal mechanisms through which an exposure variable (X) affects an outcome variable (Y) through a mediator variable (M). It decomposes the total effect of X on Y into direct and indirect effects where direct effect measures the impact of X on Y, bypassing M, while the indirect effect quantifies how X influences Y through M. Mediation analysis facilitates the understanding of the underlying mechanism by identifying and quantifying the mediation pathway, providing insights into how changes in the exposure variable translate to changes in the outcome variable through the mediator. In the case of TOTXVI, mediation analysis can help us identify particular brain connections that are both damaged by treatment (the *α* path in Figure 1a) and influence neurocognitive outcomes (the *β* path in Figure 1a), thus serving as “middle men” risk factors that foster and enable this negative pathway.

Such mediation analysis, however, requires careful modeling of the complex, high-dimensional brain connectivity network data that serve as the high-dimensional mediator. In the TOTXVI study, the structural brain connectivity network of a patient is represented by a 379 × 379 positive continuous-valued symmetric adjacency matrix, also known as the connectivity matrix. The rows and columns in the connectivity matrix correspond to brain regions of interest (ROIs), i.e. nodes. While the value of an entry at the *j*th row and the *k*th column describes the extent to which the *j*th ROI and the *k*th ROI are connected, i.e. edge weights, with larger values indicating better connectivity. Figure 1b shows a processed connectivity matrix of a randomly sampled patient in the TOTXVI real data analysis in Section 6.

One key statistical challenge in dealing with brain connectivity matrices is their high dimensionality. Specifically, recent imaging techniques are capable of measuring a large number of ROIs (e.g. *V* =379 in TOTXVI) and distinct brain connections 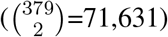 that easily exceed the sample size (Glasser et al., 2016). However, the majority of these connections are unlikely to be influenced by IT chemotherapy or exhibit effects on neurocognitive outcomes, with only a small proportion of the connections are expected to display significant nonzero effects. Additionally, the brain connectivity network displays important structural dependency (Park and Friston, 2013), with functionally important ROIs densely connected to each other forming relatively isolated modulars (Arnatkeviciute et al., 2021), with each module (a.k.a. subnetwork) specializing in specific functions (Park and Friston, 2013). Therefore, it is critical to account for both the sparsity of their effects and the unique structural characteristics of the brain connectivity networks.

Unfortunately, existing mediation analysis methods do not account for these important features associated with the complex, high-dimensional brain connectivity data. Specifically, standard mediation analysis is univariate in nature and examines mediators one at a time in the presence of high-dimensional mediators. However, such univariate mediation analysis does not account for the structural interdependence among brain connectivity and can result in low power in detecting causal mediators. To remedy the shortcoming of univariate mediation analysis, several high-dimensional mediation models have been recently proposed, motivated by a wide variety of modern data types including microbiome (Sohn and Li, 2019), gene expression (van Kesteren and Oberski, 2019; Song et al., 2020, 2021a,b), functional magnetic resonance imaging (MRI) (Chén et al., 2018; Zhao, Lindquist and Caffo, 2020), epigenetic (Zhang et al., 2016), and imaging genetics data (Bi et al., 2017). However, the existing high-dimensional mediation models are not suitable for analyzing our TOTXIV data as most of them focus on unstructured mediators and are not designed to handle brain connectivity data with structural dependency. Additionally, most high-dimensional mediation methods are designed for handling multivariate mediators with dimensions up to only a few hundred (Zhao, Li and Caffo, 2020), while the brain connectivity network in the TOTXVI study consists of tens of thousands of distinct connections. Indeed, existing methods and software packages struggle with computational speed and memory issues when applied to our data.

Here, we developed a novel scalable high-dimensional mediation analysis method with a network mediation structural shrinkage (NMSS) prior designed for accommodating network mediators such as brain connectivity. The NMSS prior imposes global sparsity to encourage the selection of a small set of connections as potentially causal mediators. It incorporates the effective degree of each node in the connectivity network to account for the network dependency structures. It also encourages the detection of subnetworks with similar effects by modeling dependencies within subnetworks, thus facilitating the detection of interpretable subnetworks and aiding the interpretation of the estimated causal effects. Importantly, NMSS also enhances the statistical power for detecting network connections with nonzero mediation effects by allowing information borrowing between the X-M and M-Y pathways. The proposed framework is compatible with observational or nonrandomized studies by allowing confounding adjustments. The method is also highly scalable to large connectivity networks with tens of thousands of edges.

The paper is organized as follows. Section 2 introduces the proposed high-dimensional mediation analysis framework for a brain connectivity mediator, the proposed NMSS prior, and the corresponding modeling assumptions. Section 3 describes the estimation and inference procedure accompanied by the algorithm. Section 4 presents simulation studies designed to evaluate the performance of the proposed method. Section 5 presents the application of the proposed method to the motivating TOTXVI clinical trial data. Section 6 concludes the paper with discussions.

## 2 MODEL

### 2.1. Model specification

Let *x*_*i*_ be a univariate exposure, *y*_*i*_ be a univariate outcome, and **m**_*i*_ be a high-dimensional mediator represented by the connectivity matrix, all for the *i*th subject (*i*∈ {1, …, *n*}). The connectivity matrix 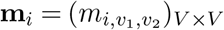 is a symmetric weighted adjacency matrix with zero diagonal elements, corresponding to an undirected network without self-loop, where self-loop represents a connection that connects a node to itself. Let *c*_1*i*_ be the exposure-mediator confounder, *c*_2*i*_ be the exposure-outcome confounder, and *c*_3*i*_ be the mediator-outcome confounder. These confounders *c*_1*i*_, *c*_2*i*_, and *c*_3*i*_ are not necessarily exclusive. To simplify the narrative, we assume these confounders to be univariate for the purpose of this presentation. However, extending the analysis to multivariate confounders is straightforward. In our motivating TOTXVI dataset, **m**_*i*_ represents a high-dimensional structural brain connectivity matrix. Each element 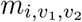 is a continuous value, with larger values indicating stronger connectivity between ROIs *v*_1_ and *v*_2_. Our goal is to identify influential elements or connections within **m**_*i*_ that mediate the treatment effect of *x*_*i*_ on the neurocognitive outcome *y*_*i*_, with adjustment of the confounders *c*_1*i*_, *c*_2*i*_, and *c*_3*i*_ (Figure 2).

**Fig 2:**
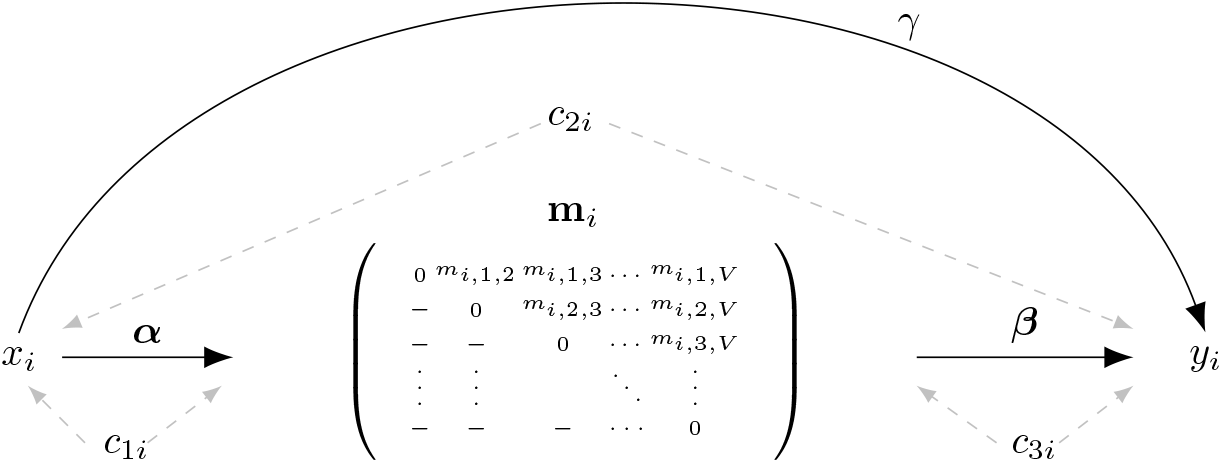
Mediation model with an exposure *x*_*i*_, a high-dimensional connectivity matrix mediator **m**_*i*_, and an outcome *y*_*i*_ as well as exposure-mediator confounder *c*_1*i*_, exposure-outcome confounder *c*_2*i*_, and mediator-outcome confounder *c*_3*i*_.

Mediation analysis aims to delineate the relations among the exposure, mediator, and outcome variables based on solving two linear equations (MacKinnon, 2008). Following this framework, we first model the relationship between the connectivity matrix mediator **m**_*i*_ and the exposure *x*_*i*_ with a matrix-on-scalar regression

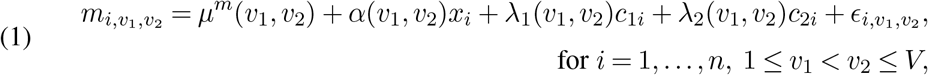

where *µ*^*m*^(*v*_1_, *v*_2_), *α*(*v*_1_, *v*_2_), *λ*_1_(*v*_1_, *v*_2_), and *λ*_2_(*v*_1_, *v*_2_) are matrix coefficients corresponding to an intercept, *x*_*i*_, *c*_1*i*_, and *c*_2*i*_, respectively. 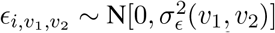 is the measurement error with unknown variance 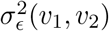. For simplicity, we let 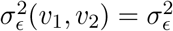 for all the numerical experiments in the paper. We only include in the equation the upper triangular elements in **m**_*i*_, indexed by 1 ≤*v*_1_ *< v*_2_ ≤ *V*, due to the symmetry of the matrix.

Next, we model the relationship between the outcome *y*_*i*_ and the connectivity matrix mediator **m**_*i*_ with a scalar-on-matrix regression

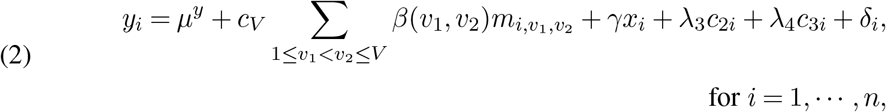

where *µ*^*y*^ is a scalar intercept, *β*(*v*_1_, *v*_2_) is the matrix coefficient corresponding to 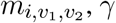 *λ*_3_ and *λ*_4_ are scalar coefficients, and *δ*_*i*_ is the measurement error following 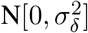. *c*_*V*_ is a normalizing constant suggested to be *O*(*V* ^−1^) for high-dimensional predictors (Kang, Reich and Staicu, 2018) and can be absorbed into the connectivity predictors 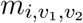 as part of the standardization procedure. Similarly, we only include in the equation the upper triangular elements of **m**_*i*_ due to the symmetry of **m**_*i*_.

### 2.2. Model assumptions and causal identifications

To define the causal effects of interest, we introduce the potential outcome notation (Rubin, 1974) corresponding to the variables in the proposed model. Specifically, let *Y*_*i*_(*x*) (corresponding to *y*_*i*_) denote the potential outcome under *X*_*i*_ = *x*, referring to the value of the outcome random variable that would have been observed had exposure *X*_*i*_ been set to *x*. Let 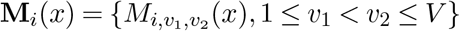 and 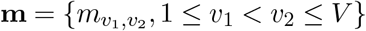 be a realization of **M**_*i*_(*x*); **M**_*i*_(*x*) under *X*_*i*_ = *x* and *Y*_*i*_(*x*, **m**) under *X*_*i*_ = *x* and **M**_*i*_ = **m** are defined similarly. We let **C**_*i*_ = {*C*_1*i*_, *C*_2*i*_, *C*_3*i*_}and **c** ={*c*_1_, *c*_2_, *c*_3_} be the variable and realization corresponding to the set of all confounders. We make the generally applied composition constraint by letting *Y*_*i*_(*x*) = *Y*_*i*_(*x*, **M**_*i*_(*x*)) (Pearl, 2000). Suppose we focus on two arbitrary levels of the exposure, *X*_*i*_ = *x* and *X*_*i*_ = *x*^*^. Causal effects are formally defined with respect to potential outcomes, which are hypothetical variables and may not be observed in real data. The identification and estimation of causal effects with observed data requires a set of assumptions. Throughout, we make the commonly used stable unit treatment value assumption (SUTVA) (Rubin, 1980). SUTVA assumes that there is no interference between subjects, implying that one individual’s exposure assignment does not affect the outcomes of others. It also assumes consistency, which states that the variables whichever observed are identical to the potential variables under the actually observed exposure or mediator values (VanderWeele, 2015). Besides SUTVA, we make the widely used no-unmeasured-confounding assumptions (VanderWeele, 2015), including

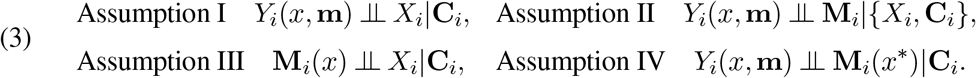

The interpretations associated with the above no-unmeasured-confounding assumptions are: I, no-unmeasured-confounding of the exposure-outcome relationship; II, no-unmeasured-confounding of the mediator-outcome relationship; III, no-unmeasured-confounding of the exposure-mediator relationship; IV, no mediator-outcome confounder that is affected by the exposure (VanderWeele and Vansteelandt, 2009). As with all causal mediation analyses, the exposure, mediators, and outcome should also satisfy the temporal ordering assumption where the exposure precedes the mediators, and the mediators precede the outcome.

Under the above assumptions, the average natural direct effect (DE), the average natural indirect effect (IE), as well as the total effect (TE) are identifiable (VanderWeele and Vansteelandt, 2014), which are obtained based on observed data via the equations presented below (Equations (4)). Particularly, TE can be decomposed into DE and IE, with IE representing the exposure effect mediated through the set of mediators. The proportion mediated (PM), defined as PM = IE/TE, is often used to assess the extent to which the total effect of the exposure on the outcome operates through the mediator. Under the proposed framework, we refer to the IE as the causal mediation effect.

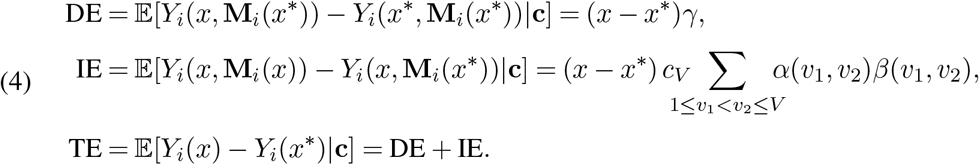

## 3. NMSS PRIOR

### 3.1. Overview

In the above model, our key parameters of interest are ***α*** = {*α*(*v*_1_, *v*_2_), 1 ≤ *v*_1_ *< v*_2_ ≤ *V*, ***β*** = *β*(*v*_1_, *v*_2_), 1 ≤ *v*_1_ *< v*_2_ ≤ *V*. We let ***θ*** = {***α, β***} = {*θ*(*ξ, v*_1_, *v*_2_), *ξ* = *α, β*, 1 ≤ *v*_1_ *< v*_2_ ≤ *V*}. For model inference, we adopt a Bayesian approach and developed a novel network mediation structural shrinkage (NMSS) prior for ***θ***, which encompasses four desirable modeling features. First, NMSS effectively induces sparsity on the matrix coefficients (*α*(*v*_1_, *v*_2_) and *β*(*v*_1_, *v*_2_)), which addresses global sparsity in the matrices and allows us to select a sparse set of connections with nonzero effects. Second, NMSS explicitly models the probability of nonzero effect as a function of the effective degree of each node, thus accounting for the dense connection among the important nodes (ROIs). Third, NMSS allows for information borrowing across the X-M and M-Y pathways to increase the power of detecting connections with mediation effects. Fourth, NMSS models the dependencies within subnetworks to promote the detection of interpretable subnetworks. The details for the NMSS prior along with its four characteristics are described below.

### 3.2. Global sparsity, the effective degree of nodes, and the detection of mediators

To facilitate the selection of nonzero mediation effects, we introduce a set of latent binary indicator variables **I** = {*I*(*ξ, v*_1_, *v*_2_), *ξ* = *α, β*, 1 ≤ *v*_1_ *< v*_2_ ≤ *V*} underlying ***θ***, such that *θ*(*ξ, v*_1_, *v*_2_) = 0 if *I*(*ξ, v*_1_, *v*_2_) = 0 and *θ*(*ξ, v*_1_, *v*_2_) ≠ 0 if *I*(*ξ, v*_1_, *v*_2_) = 1. We first specify prior for the latent binary indicator variables **I** in this subsection and then specify priors for the conditional distribution ***θ***|**I** in the next subsection.

For **I**, we propose the following prior based on the Ising distribution

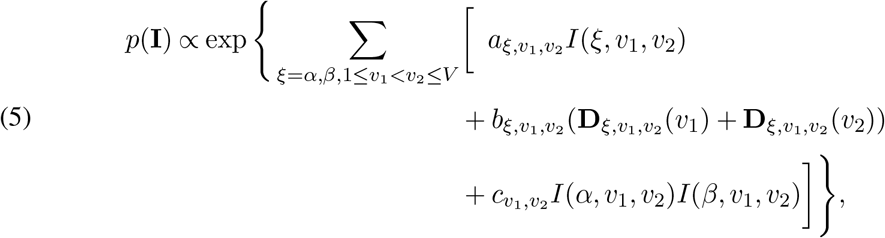

where 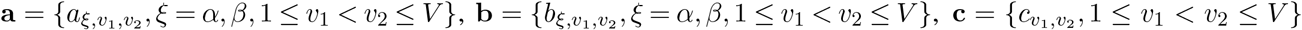 are hyperparameters fixed *a priori*; and 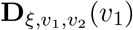 and 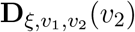 are the effective degree of node *v*_1_ and *v*_2_ with respective to the connection between *v*_1_ and *v*_2_ calculated with *I*(*ξ*, ·, ·).For the hyperparameters, we let

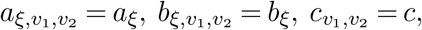

in the numerical analysis with *a*_*ξ*_ *<* 0, *b*_*ξ*_ *>* 0, and *c >* 0, for *ξ* = *α, β*. We treat *a*_*ξ*_, *b*_*ξ*_, *c* as tuning parameters of which the optimal values are chosen based on cross-validation (details below). For the effective degree of the nodes, we have

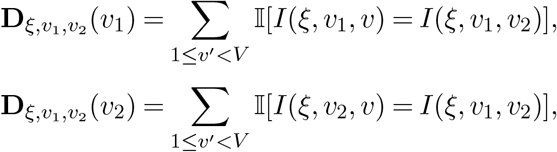

where I(·) is a characteristic function that equals to 1 if the input holds and 0 otherwise. Under the definition, for *I*(*ξ, v*_1_, *v*_2_) = 1, 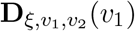 and 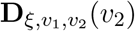 are the numbers of connections with nonzero effects connected to *v*_1_ and *v*_2_ respectively.

The role of these hyperparameters and the effective degree of the nodes can be readily understood through the following conditional prior, which is derived from the above Ising prior:

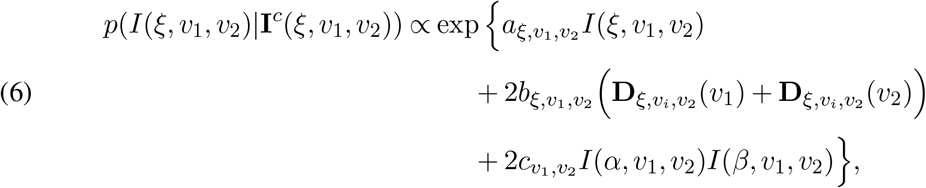

where *ξ* = *α, β*, 1 *< v*_1_ *< v*_2_ *< V* and **I**^*c*^(*ξ, v*_1_, *v*_2_) = **I \** *I*(*ξ, v*_1_, *v*_2_).

From Equation (6), the conditional probability of *I*(*ξ, v*_1_, *v*_2_) = 1|· is negatively impacted by *a*_*ξ*_ (*a*_*ξ*_ *<* 0), which effectively controls the proportion of 1s in **I** and hence the global sparsity of ***θ***. In addition, the conditional probability of *I*(*ξ, v*_1_, *v*_2_) = 1|· is positively impacted by the effective degree of node 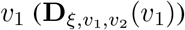 plus the effective degree of node 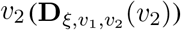, Figure 3) as controlled by *b*_*ξ*_ (*b*_*ξ*_ *>* 0). Consequently, the connection between *v*_1_ and *v*_2_ are more likely to display nonzero effect and become influential (*I*(*ξ, v*_1_, *v*_2_) = 1) if *v*_1_ and *v*_2_ are connected to more influential edges. Finally, the hyperparameter *c* (*c >* 0) effectively controls the probability of concurrence between *I*(*α, v*_1_, *v*_2_) = 1 and *I*(*β, v*_1_, *v*_2_) = 1 and thus allows for information borrowing between the X-M and M-Y pathways to facilitate the detection of edges with mediation effect with both *I*(*α, v*_1_, *v*_2_) = 1 and *I*(*β, v*_1_, *v*_2_) = 1.

**Fig 3:**
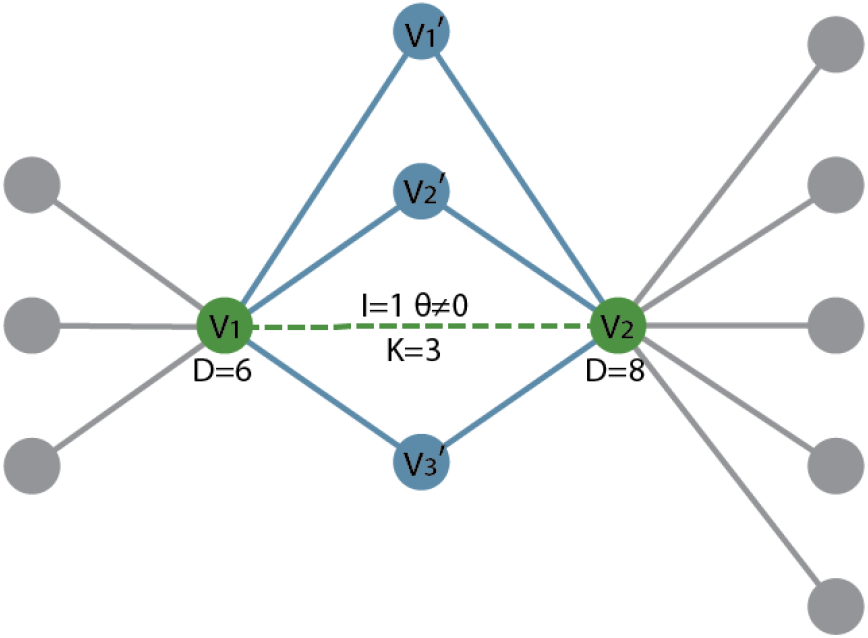
A schematic for the NMSS prior. The circles represent nodes and the lines between nodes represent nonzero coefficient of the corresponding connections. We focus on a nonzero effect (*θ*≠ 0 and *I* = 1) of the connection between node *v*_1_ and *v*_2_. As shown in the schematic, the effective degree of *v*_1_ is 6 and that of *v*_2_ is 8. There are three nodes (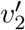 and 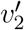) and each forms a triangular clique jointly with *v*_1_ and *v*_2_ yielded 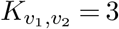. The prior mean for *θ* is the average effect size of the coefficients corresponding to the six blue edges.

### 3.3. Interpretable subnetworks

In this subsection, we provide a detailed description of the prior distribution of ***θ*** conditional on **I**. Given the binary variable selection indicator, a portion of the elements in ***θ*** are assigned to zeros, which correspond to spurious brain connections. That is, *p*(*θ*(*ξ, v*_1_, *v*_2_) | **I**(*ξ, v*_1_, *v*_2_) = 0) ∼ *δ*(0), where *δ*(0) is the Dirac delta function. We focus on the nonzero coefficients *θ*(*ξ, v*_1_, *v*_2_) corresponding to *I*(*ξ, v*_1_, *v*_2_) = 1 and design a shrinkage prior to promote the formation of interpretable subnetworks. Specifically,

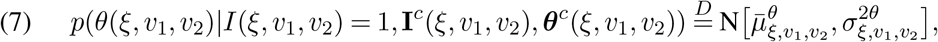

where 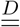 represent “distributed as”, *N* [·, ·] represent a Gaussian distribution, ***θ***^*c*^(*ξ, v*_1_, *v*_2_) = ***θ***\*θ*(*ξ, v*_1_, *v*_2_), and

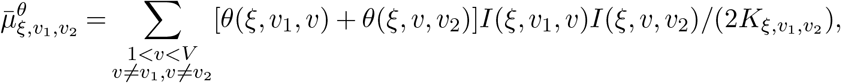

where 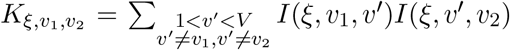 is the number of nodes with both *I*(*ξ, v*_1_, ·) = 1 and *I*(*ξ*, ·, *v*_2_) = 1. Intuitively, the prior mean 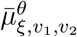 is the average value of the nonzero coefficients corresponding to the edges that form “triangular clique” together with the edge between node *v*_1_ and node *v*_2_ (Figure 3). Consequently, the prior imposes smoothness of the effects across edges within subnetworks. In the numerical analysis, we simplify the choice of hyperparameters by letting 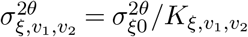. In practice, if 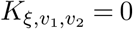 we let 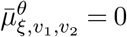 and 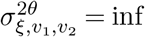 (set to a large positive number e.g. 1e4), corresponding to a non-informative prior.

### 3.4. Priors for the other parameters

Let ***µ***^*m*^ = {*µ*^*m*^(*v*_1_, *v*_2_), 1 ≤ *v*_1_ *< v*_2_ ≤ *V*}, ***λ***_1_ = {*λ*_1_(*v*_1_, *v*_2_), 1 ≤ *v*_1_ *< v*_2_ ≤ *V*}, ***λ***_2_ ={*λ*_2_(*v*_1_, *v*_2_), 1 ≤ *v*_1_ *< v*_2_ ≤*V*}, conditional priors are assigned to ***µ***^*m*^, ***λ***_1_, and ***λ***_1_ for addressing dependencies

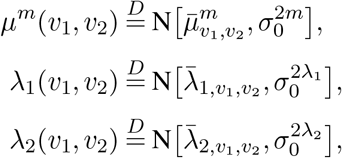

with

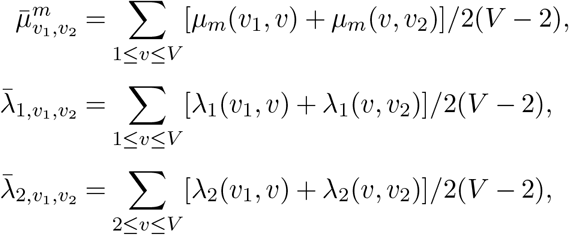

where 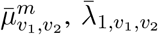 and 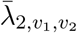 can be interpreted similarly to 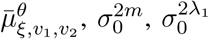, and 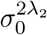 are hyperparameters that need to be fixed *a priori*.

For the rest of the unknown scalar parameters, conjugate prior distributions are assigned

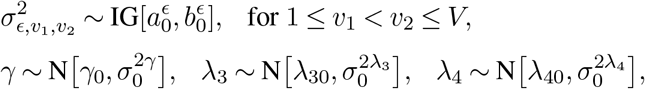

where 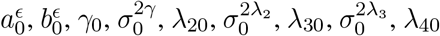 and 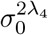 are hyperparameters fixed *a priori*.

## 4. POSTERIOR INFERENCE

### 4.1. Posterior distributions

We denote the above prior specifications, including both the prior on **I** and the conditional prior on ***θ***|**I**, as the NMSS prior. To carry out posterior inference on the model, we first denote **x** = {*x*_1_, …, *x*_*n*_}, 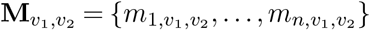, **M** = {**m**_1_, …, **m**_*n*_}, **c**_*j*_ = {*c*_*j*1_, …, *c*_*jn*_}, *j* = 1, 2, 3, ***λ***_1_ = {*λ*_1_(*v*_1_, *v*_2_), 1 ≤ *v*_1_ *< v*_2_ ≤ *V* }, ***λ***_2_ = {*λ*_2_(*v*_1_, *v*_2_), 1 ≤ *v*_1_ *< v*_2_ ≤ *V* }, and 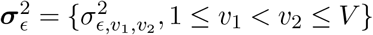. The full conditional posterior distributions of the network coefficients *θ*(*ξ, v*_1_, *v*_2_) are derived as follows. Using the NMSS prior, we have the edge-specific joint posterior distribution of {*θ*(*α, v*_1_, *v*_2_), *I*(*α, v*_1_, *v*_2_)} as

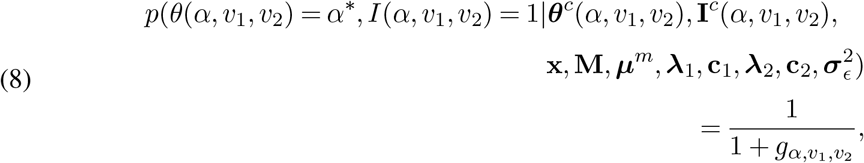

and

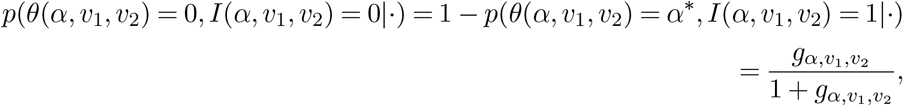

where

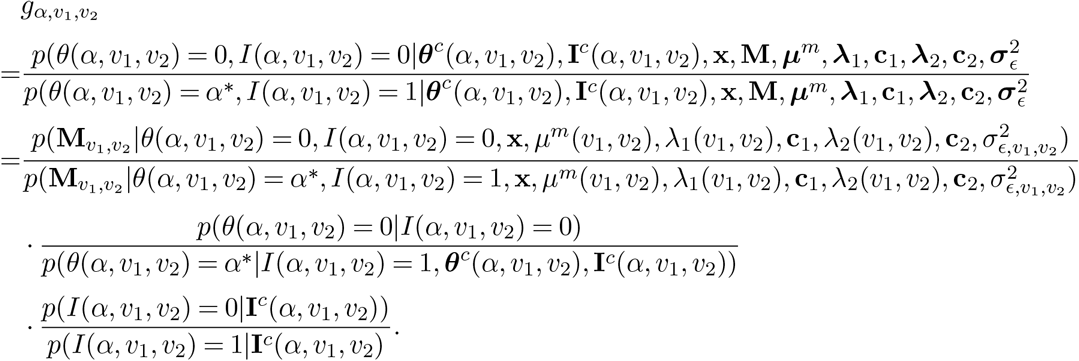

*α*^*^ is sampled from

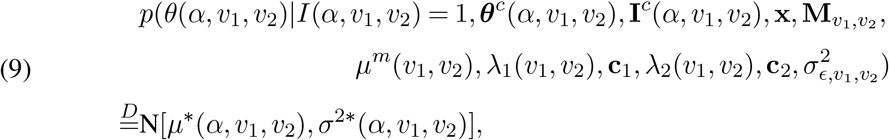

with 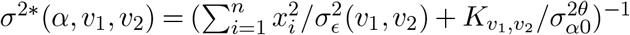 and

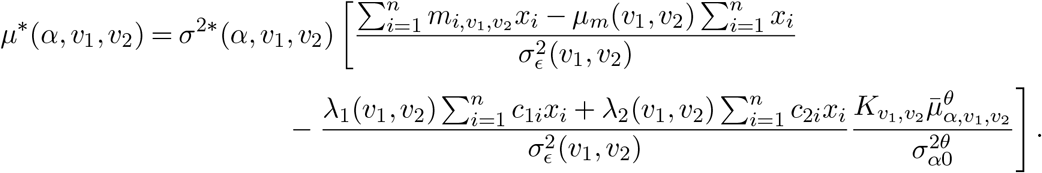

Let 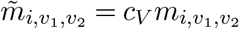, the posterior distribution of {*θ*(*β, v*_1_, *v*_2_), *I*(*β, v*_1_, *v*_2_)} can be derived similar to that of *θ*(*α, v*_1_, *v*_2_), *I*(*α, v*_1_, *v*_2_) (Supplementary Material 1). Based on Equations (6) and (7), smaller *a*_*ξ*_ (*a*_*ξ*_ *<* 0) corresponds to larger 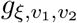 there-after smaller probability of a nonzero network coefficients i.e. smaller *p*(*θ*(*α, v*_1_, *v*_2_) = *α*^*^, *I*(*α, v*_1_, *v*_2_) = 1|·) and *p*(*θ*(*β, v*_1_, *v*_2_) = *β*^*^, *I*(*β, v*_1_, *v*_2_) = 1|·), hence encouraging the sparsity of ***α*** and ***β***. For each connection, the joint posterior distribution of the latent binary indicator and matrix coefficient {*θ*(*ξ, v*_1_, *v*_2_), *I*(*ξ, v*_1_, *v*_2_)} determines a binary choice between {*θ*(*ξ, v*_1_, *v*_2_) = *θ*^*^, *I*(*ξ, v*_1_, *v*_2_) = 1} and {*θ*(*ξ, v*_1_, *v*_2_) = 0, *I*(*ξ, v*_1_, *v*_2_) = 0} after accounting for both the prior information through the proposed NMSS prior and the likelihood (Equations (8) and (S1)). Given the binary indicator, the structural shrinkage prior for the nonzero network coefficients encourages smoothed edge effects within subnetworks.

Together, the NMSS prior leads to sparse nonzero coefficients, accounts for the dense connections among the important ROIs, addresses dependencies within subnetworks, and importantly, boosts the power for detecting mediation effects by encouraging the cooccurrence of *I*(*α, v*_1_, *v*_2_) = 1 and *I*(*β, v*_1_, *v*_2_) = 1. The calculation of the posteriors can be optimized and further made computationally efficient, as discussed in Section 4.2. The posteriors of the other parameters are provided in Supplementary Material 2.

### 4.2. Algorithm

We implement the following Markov chain Monte Carlo (MCMC) algorithm to acquire samples of unknown parameters from their posterior distributions. Specifically, for the matrix coefficients (***θ***) and their corresponding binary latent indicators (**I**), we use a single-site Gibbs sampler to generate samples from their posterior distributions by using the connection-specific probability in Equations (8) and (S1). For each connection, {*θ*(*ξ, v*_1_, *v*_2_), *I*(*ξ, v*_1_, *v*_2_)} is sampled from a Bernoulli distribution with the binary choices {0, 0} and {*θ*^*^, 1}, where *θ*^*^ (*θ*=*α* or *β*) is sampled from the distribution in Equations (9) and (S2).

#### Algorithm 1

MCMC algorithm for the network mediation model with NMSS prior

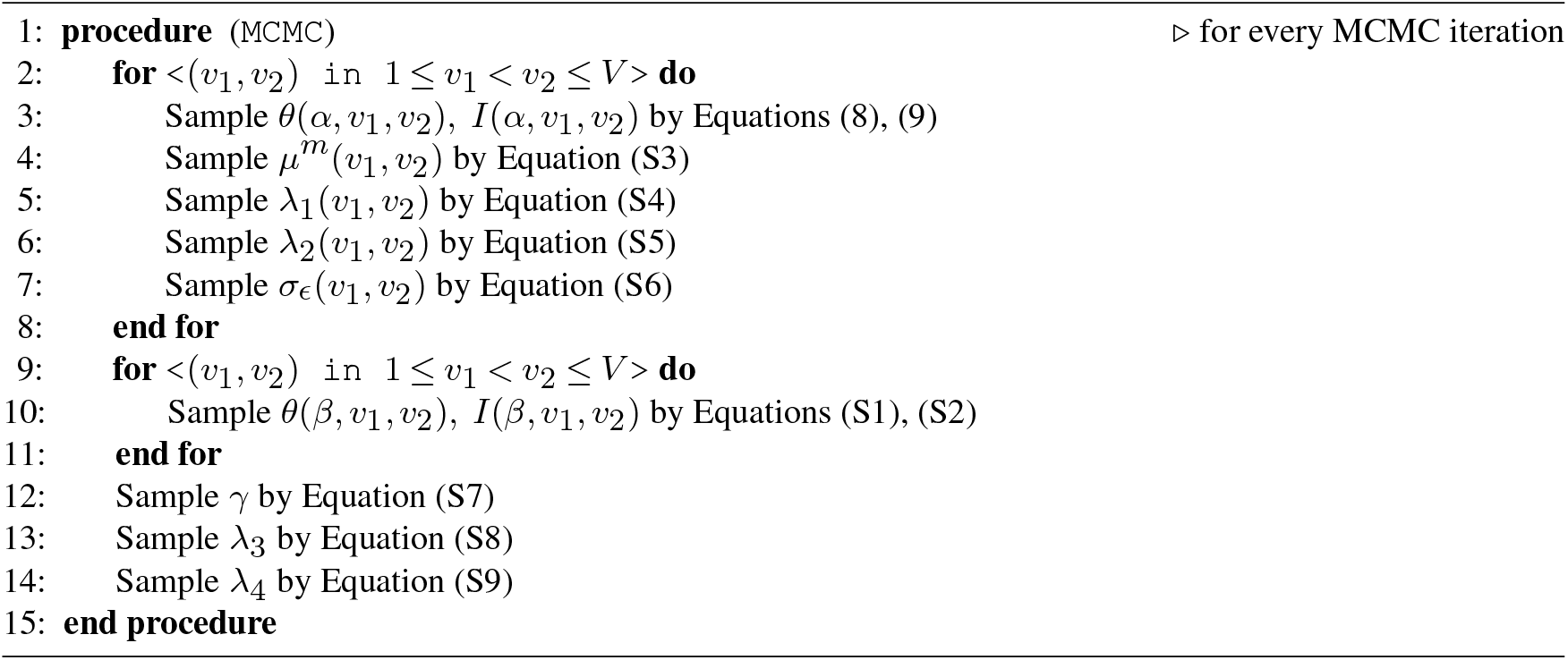

Given that *V >> n*, the computation time primarily depends on sampling the parameters indexed by *v*_1_ and *v*_2_. A naive implementation of the algorithm results in a computational complexity of *O*(*nV* ^4^) per MCMC iteration. We have developed a scalable algorithm with further optimizations to bring the computational complexity down to *O*(*nV* ^2^). Specifically, we focus on an arbitrary connection between 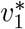 and 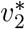 with a current value 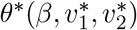 and a proposed value *β*^*^. The computational cost of 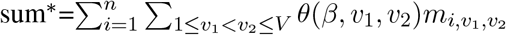 under 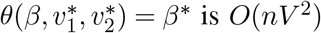 is *O*(*nV* ^2^), which, however, can be substantially reduced to *O*(*n*) with the decomposition 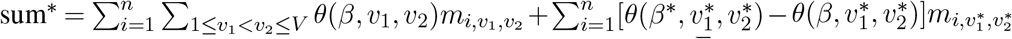. Hence, the computation complexity of updating the parameters in Equation (S1) is reduced from *O*(*nV* ^4^) to *O*(*nV* ^2^). The computation complexity of updating the parameters in Equation (8) is *O*(*nV* ^2^) based on the derived full conditional distributions. Importantly, the data related summations including 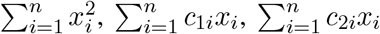 and 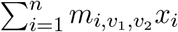 only need to be calculated once, which also saves considerable computation time. As a result, our algorithm is highly scalable.

### 4.3. Tuning parameter selection

We regard 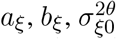, (for *ξ* = *α, β*), *c*, and 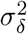 as tuning parameters of which the optimal values are chosen based on cross-validation. Specifically, *a*_*ξ*_ controls the global sparsity of *θ*(*ξ, v*_1_, *v*_2_) through *I*(*ξ, v*_1_, *v*_2_); *b*_*ξ*_ handles the importance of node through effective degrees 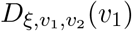 and 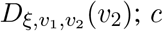 enhance the detection of mediation effect by allowing information borrowing between the X-M and M-Y pathways via addressing the cooccurrence of *I*(*α, v*_1_, *v*_2_) = 1 and 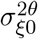 controls the smoothness of coefficients within subnetworks; and 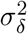 controls the under- and over-fitting of the model of Equation (2) (Goldsmith, Huang and Crainiceanu, 2014).

To select the optimal values of these tuning parameters, we implemented a cross-validation procedure (Browne, 2000) using the mean squared error of prediction as the measure-of-fit criterion (Wallach and Goffinet, 1989) with the following steps. We fix *c* = 0 to search for the optimal values of 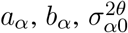 with model (1) and for the optimal values of 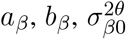, and 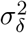 with model (2), in parallel. Afterwards, we fix 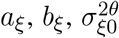, (for *ξ* = *α, β*), and 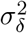 at their optimal values chosen before to perform cross-validation with both models to search for the optimal value of *c*. The detailed procedures and recommended candidate ranges of tuning parameters are provided in Supplementary Material 3.

## 5. Simulation

### 5.1. Estimation accuracy

We first conducted simulation studies to evaluate performance of the proposed NMSS method in terms of the accuracy in estimating the matrix coefficients, scalar coefficients, and causal effects. Under sample size *n* = 500, we simulated networks with *V* = 50 nodes corresponding to symmetric connectivity matrices of dimension 50 × 50. We considered different scenarios of the true matrix coefficients, which correspond to different network structures that have been observed in real brain connectivity (Park and Friston, 2013). In particular, we considered community structures (Nowicki and Snijders, 2001) where the nodes are organized into densely connected groups or communities; scale freeness (Barabási and Albert, 1999) where the degrees of nodes follows a power law with a few hub nodes of high degree and most nodes of low degree; and small-worldness (Watts and Strogatz, 1998) where nodes are more likely to be densely connected to their neighbors (Figure (S1)). We specified the true matrix coefficients ***α*** and ***β*** in the simulations based on these three network structures. Specifically, the true matrix coefficients under the community structure scenario were specified to have communities (group of nodes) of different sizes (the left panels in Figure 4 and (S1)). The true matrix coefficients under the scale freeness scenario were generated under the Barabási and Albert (Barabási and Albert, 1999) model where the degree distribution follows a power law (the middle panels in Figure 4 and (S1)). The true matrix coefficients under the small-worldness scenario were generated using the Watts and Strogatz (Watts and Strogatz, 1998) model with high local clustering and short average path lengths (the right panel in Figure 4 and (S1)).

**Fig 4:**
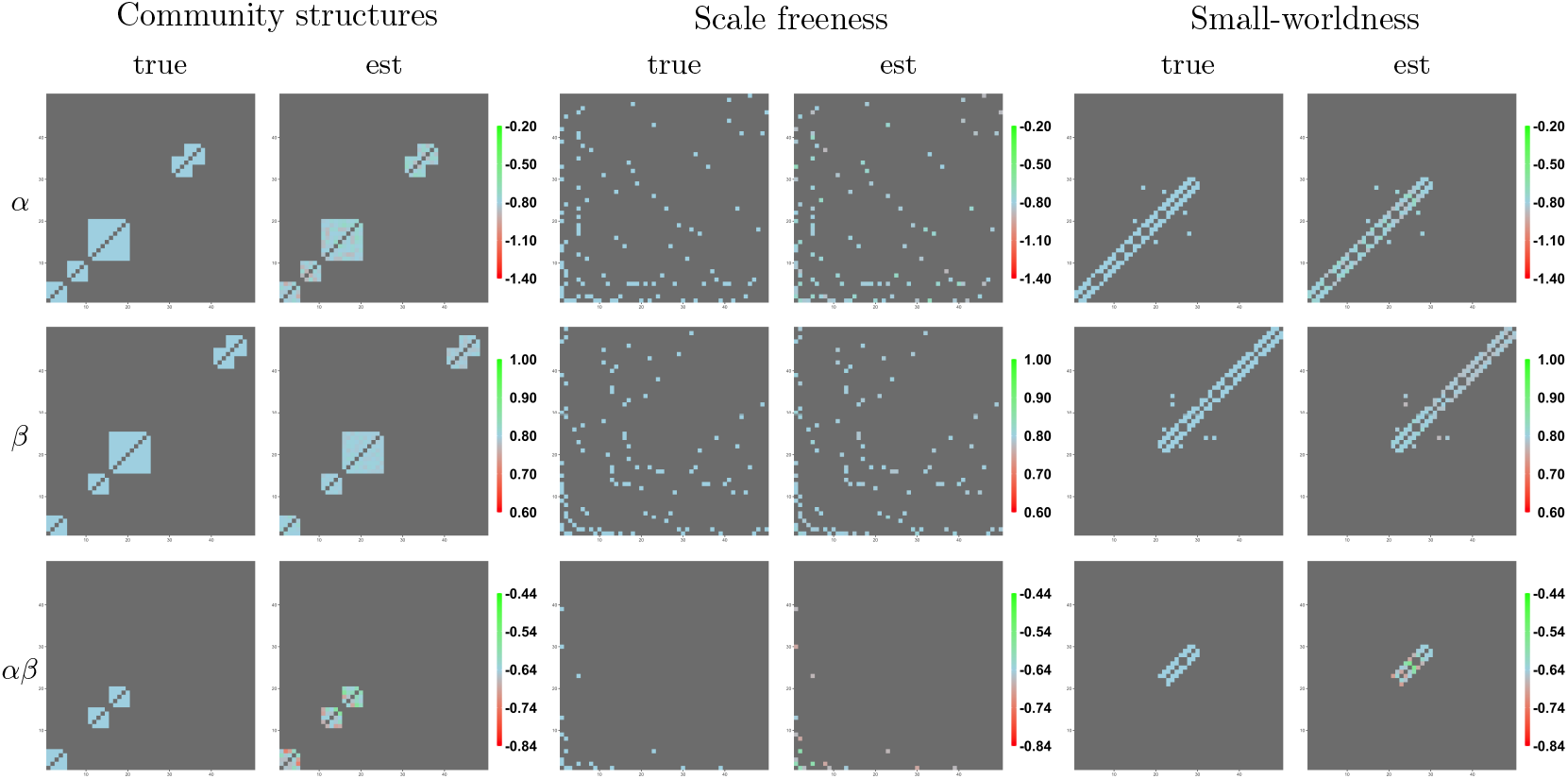
The patterns of true matrix coefficients of the simulation in Section 5.1. The 1st, 2nd, and 3rd row correspond to *α*(*v*_1_, *v*_2_), *β*(*v*_1_, *v*_2_), and *α*(*v*_1_, *v*_2_)*β*(*v*_1_, *v*_2_). Each panel, from left to right, correspond to one of the three scenarios respectively.

In each scenario, the data were simulated from Equations (1) and (2). Specifically, in Equation (1), the exposure *x*_*i*_ is a univariate continuous variable generated from a standard normal distribution. The true values of *α*(*v*_1_, *v*_2_) were generated as described above (Figure 4), where the true values of the nonzero coefficients were set to -0.8. *c*_1*i*_, *λ*_1_(*v*_1_, *v*_2_) and 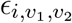 were generated from N[0, 1], N[0, 0.1^2^] and N[0, 1] respectively. For simplicity, *µ*^*m*^(*v*_1_, *v*_2_) and *λ*_2_(*v*_1_, *v*_2_) were not included. In Equation (2), the true values of *β*(*v*_1_, *v*_2_) were generated as described above (Figure 4), where the true values of the nonzero coefficients were set to 0.8. The scalar coefficients were set to *γ* = − 0.5 and *λ*_3_ = *λ*_4_ = 0.5. *δ*_*i*_ was generated from N[0, 0.1^2^]. The locations with both nonzero *α*(*v*_1_, *v*_2_) and *β*(*v*_1_, *v*_2_) correspond to the connections with nonzero mediation effects, i.e. ***α***·***β***. The true values of the causal effects are computed accordingly (Table 2).

When the proposed NMSS method is applied to the network mediation model (Section 4.2), its tuning parameters were chosen based on 5-fold cross-validation as described in Section 4.3. The proposed Algorithm (1) converged within 1,000 iterations, as suggested by several pilot chains with well-separated starting values. After 1,000 burn-in iterations, we collect the following 1,000 samples for posterior inference. The computation time for each replication is 41 minutes, using one core of an Intel(R) Xeon(R) CPU E5-2680 v2 @ 2.80 GHz. The evaluation was based on 100 replications. The estimation accuracy of the matrix coefficients was measured by the average root mean squared error (RMSE). For example, the RMSE of ***α*** with *N*_rep_ number of replications is computed as

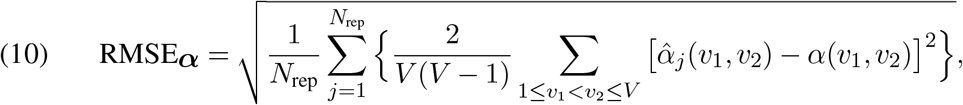

where 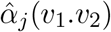 is the estimate value in the *j*th replication and *α*(*v*_1_, *v*_2_) is the corresponding true value. The RMSEs of ***β***, the element-wise product ***α***·***β***, and ***λ***_1_ are computed similarly. The summary of the matrix and scalar coefficients estimates are shown in Table 1. The summary of the causal effects estimates are shown in Table 2. The Figure 4 shows the visualization of one randomly selected replication corresponding to the three scenarios, respectively. The discrepancies in both tables are all reasonably small, indicating favorable accuracy of the proposed NMSS method across various scenarios.

**Table 1.** Estimation accuracy of the matrix coefficients (upper panel) and scalar coefficients (lower panel) of the simulation studies in Section 5.1. The discrepancy of a matrix coefficient is measured by RMSE (Equation (10)); the discrepancy of a scalar coefficient is measured by bias (RMSE).

| RMSE | $\alpha$ | $\beta$ | $\alpha \cdot \beta$ | $\lambda_1$ |
| --- | --- | --- | --- | --- |
| community structures | 0.0117 | 0.0015 | 0.0058 | 0.0440 |
| scale freeness | 0.0090 | 0.0013 | 0.0031 | 0.0439 |
| small-worldness | 0.0094 | 0.0025 | 0.0042 | 0.0439 |
| bias (RMSE) | $\gamma$ | $\lambda_3$ | $\lambda_4$ | |
| community structures | 0.0131 (0.0443) | 0.000 (0.0047) | 0.0003 (0.0050) |  |
| scale freeness | 0.0077 (0.0200) | -0.0001 (0.0045) | -0.0002 (0.0049) |  |
| small-worldness | 0.0072 (0.0291) | 0.0006 (0.0058) | -0.0006 (0.0056) |  |

**Table 2.** Estimation accuracy of the causal effects of the simulation studies in Section 5.1. The discrepancies of DE, IE, and TE are measured by bias (RMSE)

|  | DE |  | IE |  | TE |  |
| --- | --- | --- | --- | --- | --- | --- |
|  | true | bias (RMSE) | true | bias (RMSE) | true | bias (RMSE) |
| community structures | -0.5 | 0.0131 (0.0443) | -19.2 | 0.0544 (0.2231) | -19.7 | 0.0675 (0.2163) |
| scale freeness | -0.5 | 0.0077 (0.0200) | -5.76 | 0.0132 (0.1191) | -6.26 | 0.0209 (0.1168) |
| small-worldness | -0.5 | 0.0072 (0.0291) | -9.60 | 0.0341 (0.1335) | -10.1 | 0.0413 (0.1263) |

### 5.2. Competing methods

#### 5.2.1. Estimation accuracy

We compared the proposed NMSS method with a recently developed pathway mediation analysis (PMA) model (Zhao, Li and Caffo, 2020). PMA introduces a penalized optimization approach for pathway analysis in multimodal neuroimaging data integration. It allows up to two data modalities to serve as separate sets of mediators, such as structural and functional connectivity, and uses a Lasso-type penalty to induce sparse estimates of pathway effects, aiming to reveal the roles of mediators in the treatment-outcome pathway by defining and estimating pathway effects. PMA does not allow for confounders and requires the exposure to be either binary or categorical. It proceeds with vectorized connectivity mediators, treating all connections of a brain network as fully exchangeable. Notably, it cannot handle ultra-high dimensional mediators due to its heavy computational cost. For the present study, we considered the single-modality PMA where the pathway effect *α*(*v*_1_, *v*_2_)*β*(*v*_1_, *v*_2_) is analogous to the mediation effect. PMA involves a number of tuning parameters, namely {*ν*_1_, *ν*_2_, *ρ, κ*_1_, *κ*_2_, *κ*_3_, *κ*_4_, *µ*_1_, *µ*_2_}. Following Zhao, Li and Caffo (2020), we fix *ν*_1_ = *ν*_2_ = 2 and *ρ* = 1, set *κ*_1_ = *κ*_2_ = *κ*_3_ = *κ*_4_ =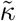, *µ*_1_ = *µ*_2_ =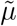, and run grid search to minimize a modified Bayesian information criterion (BIC) to obtain the optimal 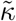 and 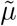 choices.

We carried out the comparison under the three scenarios similar to simulation 1 (Section 5.1) with a few adjustments due to the aforementioned limitation of PMA. These adjustments include: 1) excluded all confounders; 2) used a binary exposure *x*_*i*_ with *P* (*x*_*i*_ = 1) = 0.5; 3) based on *N*_rep_ = 20 instead of 100 to keep memory consumption and computation time. The other model setups and data generation processes were identical to simulation 1. The comparison of the estimated matrix coefficients and scalar coefficients are shown in Table 3, and the comparison of the estimated causal effects are shown in Table 4 indicating our proposed NMSS method produced more accurate estimates for both matrix coefficients and causal effects across scenarios compared to PMA. The heat maps of the estimated matrix coefficients in one randomly picked replication are visualized in Supplementary Figure (S2)- (S4).

**Table 3.** Comparison of the estimated matrix and scalar coefficients produced by NMSS and PMA.

| | | $\alpha$ | $\beta$ | $\alpha \cdot \beta$ | $\gamma$ |
| --- | --- | --- | --- | --- | --- |
|  |  | RMSE | RMSE | RMSE | bias (RMSE) |
| community structures | NMSS | 0.0169 | 0.0015 | 0.0079 | -0.0103 (0.0327) |
|  | PMA | 0.0821 | 0.0204 | 0.0187 | -1.1152 (1.1493) |
| scale freeness | NMSS | 0.0134 | 0.0013 | 0.0047 | 0.0092 (0.0170) |
|  | PMA | 0.0716 | 0.0189 | 0.0125 | -0.3454 (0.3611) |
| small-worldness | NMSS | 0.0131 | 0.0016 | 0.0055 | -0.0056 (0.0216) |
|  | PMA | 0.0863 | 0.0040 | 0.0146 | -0.1349 (0.1746) |

**Table 4.** Comparison of the estimated causal effects produced by NMSS and PMA.

|  |  | DE | IE | TE |
| --- | --- | --- | --- | --- |
|  |  | bias (RMSE) | bias (RMSE) | bias (RMSE) |
| community structures | NMSS | -0.0103 (0.0327) | 0.1896 (0.3474) | 0.1793 (0.3396) |
|  | PMA | -1.1152 (1.1493) | 1.5550 (1.6647) | 0.4399 (0.7366) |
| scale freeness | NMSS | 0.0092 (0.0170) | 0.0163 (0.1397) | 0.0255 (0.1401) |
|  | PMA | -0.3454 (0.3611) | 0.5508 (0.6651) | 0.2054 (0.4804) |
| small-worldness | NMSS | -0.0056 (0.0216) | 0.0577 (0.2261) | 0.0521 (0.2182) |
|  | PMA | -0.1349 (0.1746) | 0.2594 (0.5832) | 0.1245 (0.5649) |

#### 5.2.2. ROC

To further demonstrate the sensitivity and specificity of the proposed NMSS method in comparison with PMA, we designed a scenario with a lower signal-to-noise ratio (SNR). Specifically, we adopted similar scenarios as the previous simulation (Section 5.2.1) but instead reducing the true effect sizes of the nonzero matrix coefficients to *α*(*v*_1_, *v*_2_) = −0.1 and *β*(*v*_2_, *v*_2_) = 0.05, yielding *α*(*v*_1_, *v*_2_)*β*(*v*_2_, *v*_2_) =−0.005. In addition to regular NMSS, we also considered a degenerated version of NMSS denoted by NMSSD, where the hyperparameter *c* (or 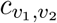 in Equation (5) is fixed at 0. Hence, *I*(*α, v*_1_, *v*_2_) and *I*(*β, v*_1_, *v*_2_) become independent and there is no information borrowing between X-M and Y-M pathways anymore in NMSS-D. We compare the receiver operating characteristic (ROC) curves across the three methods. The ROCs under the community structures scenario are shown in Figure 5. Compared to PMA, both NMSS and NMSS-D achieved superior performance in *α*(*v*_1_, *v*_2_), *β*(*v*_1_, *v*_2_), and *α*(*v*_1_, *v*_2_)*β*(*v*_1_, *v*_2_) demonstrating the advantage of the proposed framework. Moreover, NMSS outperforms the other two methods especially in term of ***α*** · ***β***, indicating the information borrowing between the X-M and Y-M pathways is beneficial. Intuitively, NMSS encourages *I*(*β, v*_1_, *v*_2_) = 1 and nonzero *β*(*v*_1_, *v*_2_) on connections with *I*(*α, v*_1_, *v*_2_) = 1 and nonzero *α*(*v*_1_, *v*_2_), and vice versa. Hence improving the power of detecting mediator connections characterized by both nonzero *α*(*v*_1_, *v*_2_) and *β*(*v*_1_, *v*_2_). The ROCs corresponding to the other two scenarios are shown in Supementary Figures (S5) and (S6), where NMSS outperforms PMA under the scale freeness scenario and performs comparatively under the samll-worldness scenario.

**Fig 5:**
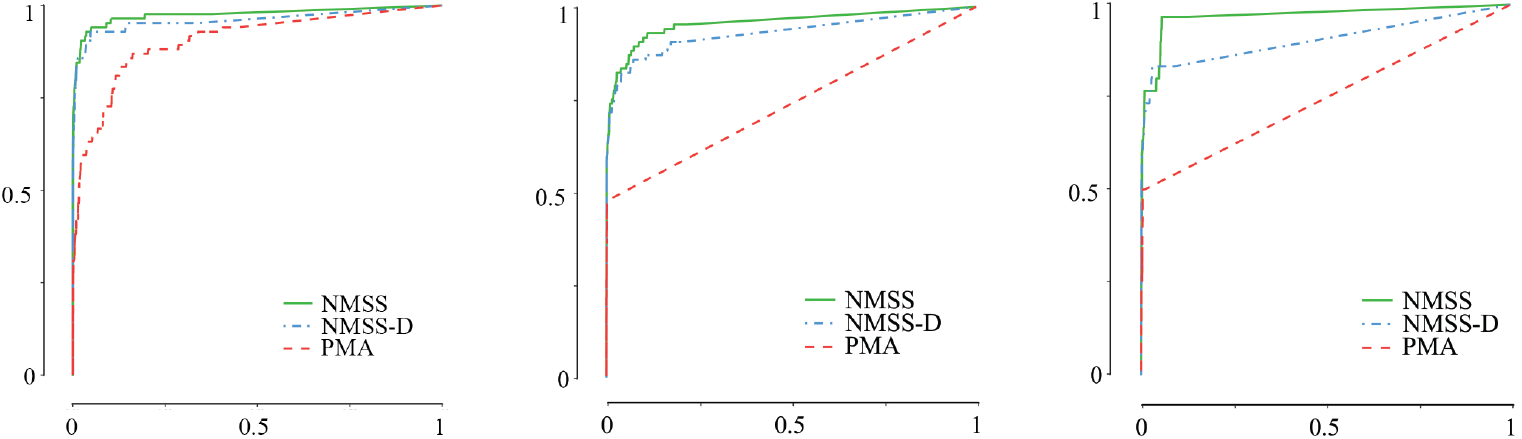
Comparison of ROC curves under low SNR in the community structures scenario across NMSS, BMSS-D, and PMA methods.

## 6. Real data analysis

### 6.1. Description

We applied our proposed NMSS method to the TOTXVI dataset to examine whether the IT-related visual processing speed deficit is mediated by brain connectivity development. Specifically, the exposure variable (*x*_*i*_) is the IT intensity, defined as the frequency of IT chemotherapy administration from the remission induction I to the end the of continuation treatment (week 120). The mediator variables (**m**_*i*_) are the brain structural connectivity development measured during the course of treatment. Such connectivity matrices were first measured by DTI combined with an anatomical parcellation of 379 brain regions (360 cortical and 19 subcortical) using the Glasser atlas (Glasser et al., 2016). For each patient, the change in brain connectivity was calculated by subtracting the connectivity matrix at remission induction I from the connectivity matrix at the end of continuation treatment (week 120). Connections 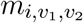 where over 25% of patients had zero values were excluded, with the corresponding variable selection indicators set at zero, i.e *I*(*α, v*_1_, *v*_2_) = 0 and *I*(*β, v*_1_, *v*_2_) = 0, yielding 25,431 remaining connections. The remaining connections were element-wise Box-Cox transformed and standardized. The neurocognitive outcome of interest (*y*_*i*_) is the visual processing speed score derived from the Woodcock-Johnson III Tests of Cognitive Abilities (Vought and Dean, 2011) measured at the end of continuation treatment (week 120) standardized by age. For the choice of the exposure-mediator confounder (*c*_1*i*_) and exposure-outcome confounder (*c*_2*i*_), given the limited sample size in this pediatric cancer study, we included age at diagnosis and biological sex, as these are known to be clinically significant factors (Bhojwani et al., 2014). We included the baseline estimated global intelligence score along with age and biological sex as the mediator-outcome confounders (*c*_3*i*_) because these factors can be associated with both visual processing speed and brain structural connectivity. Baseline intelligence, in particular, has been shown to influence both cognitive outcomes and neural connectivity (Pamplona et al., 2015). The tuning parameters were selected based on 5-fold cross-validation, and the hyperparameters were chosen to form noninformative priors with large variances (1e4). The proposed single-site Gibbs sampler converged within 1,500 iterations, as indicated by several pilot runs with well-separated starting values. After a burn-in phase of 1,500 iterations, the subsequent 2,000 samples were collected for posterior inference.

### 6.2. The influential brain connections

The parameters 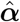 represent the effect of IT intensity (*x*_*i*_) on brain structural connectivity development (**m**_*i*_). The majority of the influential connections in 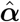 are estimated to be negative (Figure (S7)), which aligns with the anticipation that increased IT intensity is associated with decreased brain structural development. This negative correlation suggests that higher frequencies of IT administration may contribute to the deterioration of brain microstructure. We identified the top densely connected ROIs in 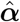 by assessing the number of influential connections for each ROI. The densely connected ROIs in 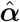 are often associated with visual movements and executive functions, such as Left Middle Temporal Area (L_MT) that involved in processing motion and visual stimuli (Table 5). We identified *α*(*v*_1_, *v*_2_) with top effect sizes and noticed that they are related to various brain regions indicating the widespread IT chemotherapy damage across the brain (Table 6).

**Table 5.** ROIs with top numbers of influential connections (#) corresponding to 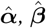, and 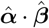.

| Rank | $\hat{\alpha}$ | | $\hat{\beta}$ | | $\hat{\alpha} \cdot \hat{\beta}$ | |
| --- | --- | --- | --- | --- | --- | --- |
|  | ROI | # | ROI | # | ROI | # |
| 1 | L_FOP2 | 27 | L_IP1 | 17 | L_IP1 | 5 |
| 2 | L_STSva | 27 | L_VMV3 | 13 | L_V3CD | 2 |
| 3 | R_PBelt | 24 | L_TGv | 12 | L_V3 | 2 |
| 4 | R_RSC | 23 | R_V3 | 11 | L_LIPv | 1 |
| 5 | L_V3 | 19 | BrStem | 10 | L_a9-46v | 1 |
| 6 | L_MT | 8 | R_Thal | 10 | L_A4 | 1 |
| 7 | L_6ma | 8 | L_24dd | 10 | R_TGd | 1 |
| 8 | L_46 | 7 | L_MT | 10 | L_47l | 1 |
| 9 | R_POS1 | 7 | L_25 | 10 | L_25 | 1 |
| 10 | L_8BM | 7 | L_PHT | 10 | R_VMV2 | 1 |

**Table 6.**
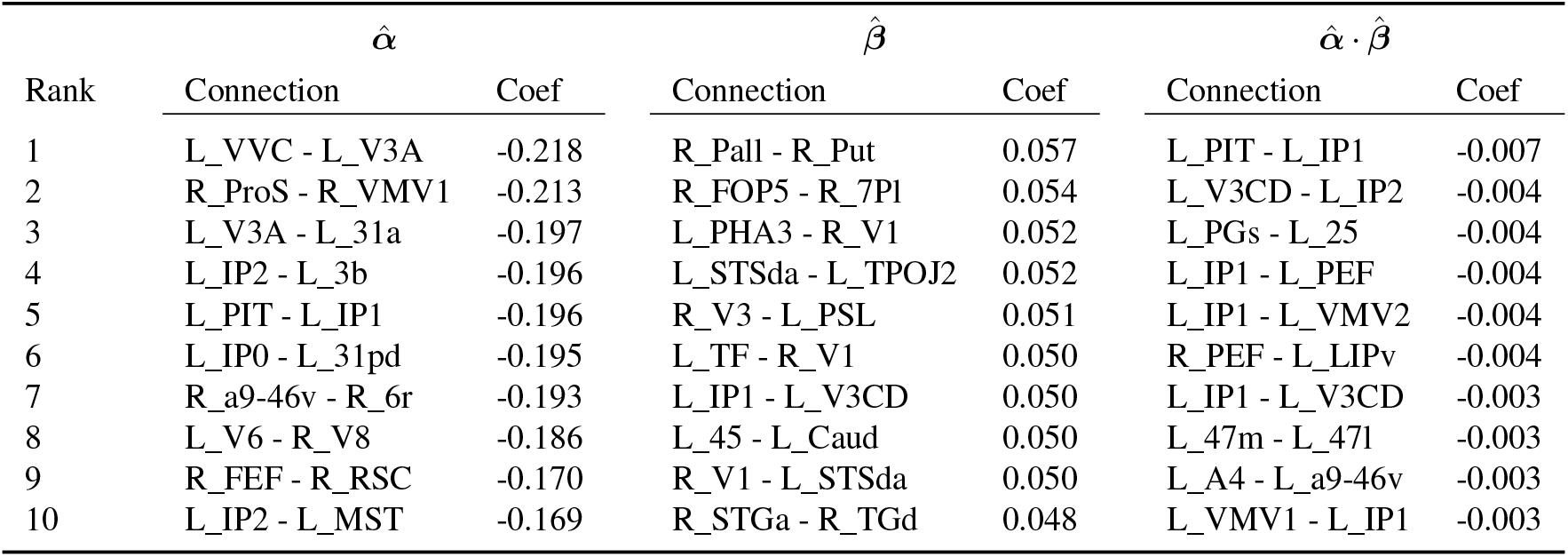
Influential coefficients with top effect sizes in 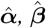, and 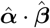.

The parameters 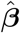 represent the effect of brain structural connectivity development (**m**_*i*_) on the visual processing speed (*y*_*i*_). Most of the significant connections in 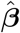 are positive (Figure (S7)), which aligns with the anticipation that better brain connectivity development is associated with favorable visual processing speed performance. This finding indicates that well-developed and efficiently connected brain network plays a critical role in maintaining high cognitive performance such as processing speed (Mash et al., 2023). The top densely connected ROIs in 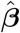 can be associated with visual, auditory, and motor processing, as well as higher-order cognitive functions such as attention, memory, and task switching. For example, one identified ROI is Left Ventromedial Visual Area 3 (L_VMV3), which is involved in identification and processing of visual stimuli associated with cognitive processing speeds (Table 5). The influential connections *β*(*v*_1_, *v*_2_) with top effect sizes are mostly related to various aspects of processing speeds (Table 6).

The product of these two sets of parameters, 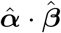, corresponds to the mediation effects of the connections, which is our primary interest. Most of the estimated nonzero effects in 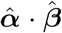 have negative signs (Figure 6), mostly arising from negative 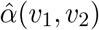 and positive 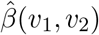. This confirms our hypothesis that brain structural connectivity development mediates the negative effect of high IT intensity on visual processing speed. In particular, the negative 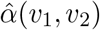 values indicate that increased IT intensity adversely affects brain connectivity development, while the positive 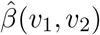 values suggest that better-developed brain connectivity is associated with favorable visual processing speed. Therefore, the negative mediation effect observed supports that intensive IT chemotherapy can lead to poorer cognitive outcomes through its detrimental impact on neural development. The top densely connected ROIs in 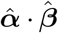 are mostly related to visual processing such as Left Intraparietal Area 1 that is involved in visuospatial attention and hand-eye coordination (Table 5). The influential connections 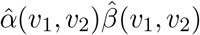 with top effect sizes often connects a visual-related ROI to an ROI with various related functions, mostly on processing speeds (Table 6). One important finding is the negative mediation effect of the connection between Right Frontal Eye Field (R_PEF) and Left Lateral Intraparietal Visual area (L_LIPv), indicating that reduced co-function between eye movement control and visuospatial attention mediates the IT-related reduced processing speed. Importantly, we identified subnetworks in 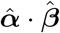 that account for the mediation effect. For example, one major subnetwork with uniformly negative mediation effects consists of a key ROI called Left Intraparietal Area 1 (L_IP1), which is involved in visuospatial attention and hand-eye coordination, together with other six visual or spatial processing related ROIs (L_VMV1, L_V3CD, L_VMV2, L_PEF, L_PIT, and L_IP2). The central role of L_IP1 in this subnetwork suggests that the region is crucial for integrating visual and spatial information, coordinating eye movements, and recognizing objects. These findings underscore the importance of monitoring and potentially adjusting IT chemotherapy such as its agents, dose, and timing, to mitigate its adverse effects on brain development and thereafter, cognitive functions.

**Fig 6:**
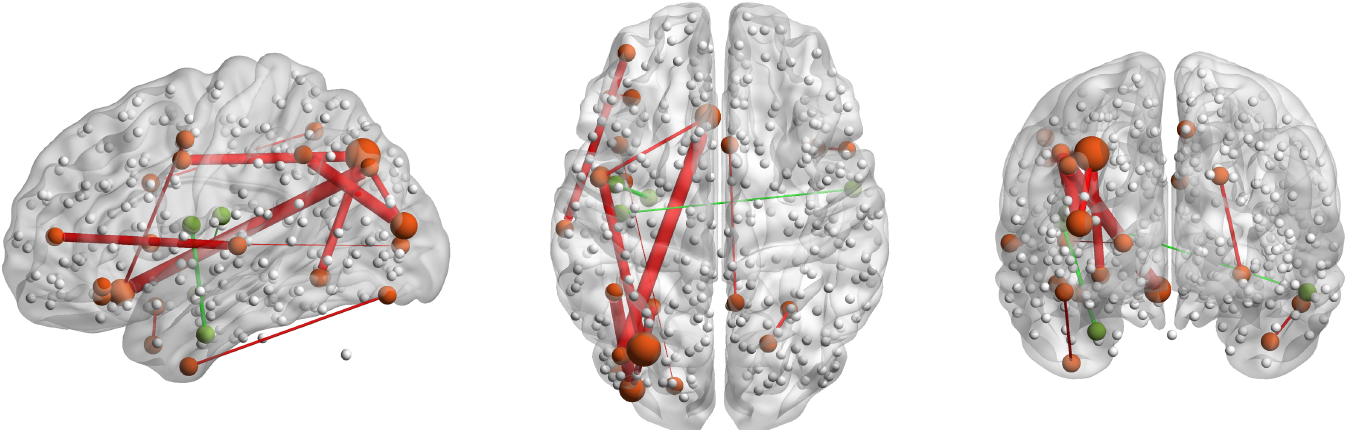
Visualization of the estimated matrix coefficients 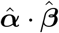 in the TOTXVI analysis.

### 6.3. The causal effects

The estimated causal effects were calculated based on Equations (4) with the estimated coefficients and the observed data. The IE, representing the mediation effect of brain structural connectivity development between IT intensity and visual processing speed, is -0.042. The DE, representing the portion of the treatment effect on visual processing speed that bypasses the brain structural connectivity development, is -0.057. The total effect (TE), which is the sum of the IE and DE, is -0.099. Therefore, the PM, calculated as IE/TE, is 42.73%. This indicates that 42.73% of the negative effect of IT intensity on visual processing speed is mediated through the development of brain structural connectivity. This substantial proportion highlights the importance of incorporating brain structural connectivity in understanding and addressing neurocognitive deficits in ALL survivors. Further research may be conducted to explore protective strategies and therapeutic interventions aimed at preserving certain brain connections during treatment or repairing connections after treatment, ultimately enhancing cognitive outcomes for pediatric ALL patients.

## 7. Discussion

In this study, we introduced the NMSS method for conducting mediation analysis with a high-dimensional brain structural connectivity mediator, addressing a critical need in understanding the neurocognitive impacts of IT chemotherapy in pediatric ALL patients. The NMSS method’s strength lies in its ability to handle high-dimensional data by incorporating global sparsity and structural dependency including the effective degree of nodes and the modularity of subnetworks. This approach enhances the detection power for identifying brain connections with mediation effects. Our findings offer important insights into how brain connectivity development mediates the adverse effects of IT intensity on visual processing speed.

Despite the promising results, there are several limitations to our study. First, while the NMSS method accounts for sparsity and structural dependency in the data, it does not explicitly model the temporal dynamics of brain connectivity changes. Incorporating temporal models could provide a more comprehensive understanding of how brain connectivity evolves over the course of treatment. Second, our analysis focused solely on visual processing speed as the cognitive outcome. Examining other neurocognitive domains, such as attention and executive function, could yield additional insights into the broader impacts of IT chemotherapy. Lastly, future work may also explore the integration of multimodal data, such as combining structural connectivity with functional connectivity and genetic data, to better understand the complex interplay between different biological systems.

In conclusion, the NMSS method provides a promising framework for analyzing highdimensional brain connectivity data in mediation analysis. Our findings with respect to the TOTXVI data contribute to a deeper understanding of the treatment-related neurocognitive impacts mediated by brain connectivity in pediatric ALL patients and offer valuable insights that could potentially aid the development of interventions to enhance the quality of life for pediatric ALL survivors.

## Acknowledgments

This study is supported by Cancer Center Support (CORE) Grant No. CA21765 and Grant No. CA90246 (W.E.R.) from the National Cancer Institute.

## Notes

### Competing Interest Statement

The authors have declared no competing interest.

