## Supplementary Material for "High-Dimensional Mediation Analysis with Network Mediators: Applications to Pediatric Acute Lymphoblastic Leukemia"

#### 1 Posteriors of $\boldsymbol{\beta}$

Similar to $\{\theta(\alpha, v_1, v_2), I(\alpha, v_1, v_2)\}$, the edge-specific joint full conditional posterior distribution of $\{\theta(\beta, v_1, v_2), I(\beta, v_1, v_2)\}$ is

$$\begin{aligned} &p(\theta(\beta, v_1, v_2) = \beta^*, I(\beta, v_1, v_2) = 1 | \boldsymbol{\theta}^c(\beta, v_1, v_2), \mathbf{I}^c(\beta, v_1, v_2), \mathbf{x}, \mathbf{M}, \gamma, \lambda_3, \mathbf{c}_3, \lambda_4, \mathbf{c}_4) \\ &= \frac{1}{1 + g_{\beta, v_1, v_2}}, \end{aligned} \tag{1}$$

where

$$\begin{aligned} &g_{\beta,v_1,v_2} \\ =&\frac{p(\theta(\beta, v_1, v_2) = 0, I(\beta, v_1, v_2) = 0 | \boldsymbol{\theta}^c(\beta, v_1, v_2), \mathbf{I}(\beta, v_1, v_2), \mathbf{x}, \mathbf{M}, \gamma, \lambda_3, \mathbf{c}_2, \lambda_4, \mathbf{c}_3)}{p(\theta(\beta, v_1, v_2) = \beta^*, I(\beta, v_1, v_2) = 1 | \boldsymbol{\theta}^c(\beta, v_1, v_2), \mathbf{I}(\beta, v_1, v_2), \mathbf{x}, \mathbf{M}, \gamma, \lambda_3, \mathbf{c}_2, \lambda_4, \mathbf{c}_3)} \\ =&\frac{p(\mathbf{y} | \theta(\beta, v_1, v_2) = 0, I(\beta, v_1, v_2) = 0, \boldsymbol{\theta}^c(\beta, v_1, v_2), \mathbf{I}^c(\beta, v_1, v_2), \mathbf{x}, \mathbf{M}, \gamma, \lambda_3, \mathbf{c}_2, \lambda_4, \mathbf{c}_3)}{p(\mathbf{y} | \theta(\beta, v_1, v_2) = \beta^*, I(\beta, v_1, v_2) = 1, \boldsymbol{\theta}^c(\beta, v_1, v_2), \mathbf{I}^c(\beta, v_1, v_2), \mathbf{x}, \mathbf{M}, \gamma, \lambda_3, \mathbf{c}_2, \lambda_4, \mathbf{c}_3)} \\ &\cdot \frac{p(\theta(\beta, v_1, v_2) = 0 | I(\beta, v_1, v_2) = 0)}{p(\theta(\beta, v_1, v_2) = \beta^* | I(\beta, v_1, v_2) = 1, \boldsymbol{\theta}^c(\beta, v_1, v_2), \mathbf{I}^c(\beta, v_1, v_2))} \\ &\cdot \frac{p(I(\beta, v_1, v_2) = 0 | \mathbf{I}^c(\beta, v_1, v_2))}{p(I(\beta, v_1, v_2) = 1 | \mathbf{I}^c(\beta, v_1, v_2))}. \end{aligned}$$

$\beta^*$ is sampled from

$$\begin{aligned} &p(\theta(\beta, v_1, v_2) | I(\beta, v_1, v_2) = 1, \boldsymbol{\theta}^c(\beta, v_1, v_2), \mathbf{I}^c(\beta, v_1, v_2), \sigma^2_\delta, \mathbf{x}, \mathbf{M}, \gamma, \lambda_3, \mathbf{c}_2, \lambda_4, \mathbf{c}_3) \\ &\stackrel{D}{=} \mathrm{N}[\mu^*_{\beta,v_1,v_2}, \sigma^{2*}_{\beta,v_1,v_2}], \end{aligned} \tag{2}$$

with $\sigma^{2*}_{\beta,v_1,v_2} = (\sum_{i=1}^n \tilde{m}^2_{i,v_1,v_2}/\sigma^2_\delta + K_{v_1,v_2}/\sigma^{2\theta}_{\beta 0})^{-1}$ and

$$\begin{aligned} \mu^*_{\beta,v_1,v_2} = \sigma^{2*}_{\beta,v_1,v_2} \Bigg\{ \frac{1}{\sigma^2_\delta} \Bigg[ &\sum_{i=1}^n (y_i - \gamma x_i - \lambda_3 c_{2i} - \lambda_4 c_{3i}) \tilde{m}_{i,v_1,v_2} \\ &- \sum_{i=1}^n \tilde{m}_{i,v_1,v_2} \sum_{(j,k)\neq(v_1,v_2)} \theta(\beta, j, k) \tilde{m}_{i,j,k} \Bigg] + \frac{K_{v_1,v_2} \bar{\mu}^\theta_{\beta,v_1,v_2}}{\sigma^{2\theta}_{\beta 0}} \Bigg\}. \end{aligned}$$

#### 2 Others posteriors

Given the prior $\mu^m(v_1, v_2) \sim \mathrm{N}[\bar{\mu}^m_{v_1,v_2}, \sigma^{2m}_0]$, the posterior distribution of $\mu^m(v_1, v_2)$ is

$$\begin{aligned} &p(\mu^m(v_1, v_2) | \boldsymbol{\mu}_{-(v_1,v_2)}, \alpha(v_1, v_2), \mathbf{x}, \mathbf{M}_{v_1,v_2}, \lambda_1(v_1, v_2), \mathbf{c}_1, \lambda_2(v_1, v_2), \mathbf{c}_2, \sigma^2_\epsilon(v_1, v_2)) \\ &\stackrel{D}{=} \mathrm{N}[\mu^{*m}_{v_1,v_2}, \sigma^{2*m}_{v_1,v_2}] \end{aligned} \tag{3}$$

where $\boldsymbol{\mu}_{-(v_1,v_2)}$ is the subset of $\boldsymbol{\mu}$ with the element corresponding to $(v_1, v_2)$ removed, and

$$\begin{aligned}\sigma^{2*m}_{v_1,v_2} =& \frac{1}{n/\sigma^2_\epsilon(v_1,v_2) + 1/\sigma^{2m}_0},\\ \mu^{*m}_{v_1,v_2} =& \left[\frac{\sum_{i=1}^n m_{i,v_1,v_2} - \alpha(v_1,v_2)\sum_{i=1}^n x_i - \lambda_1(v_1,v_2)\sum_{i=1}^n c_{1i} - \lambda_2(v_1,v_2)\sum_{i=1}^n c_{2i}}{\sigma^2_\epsilon(v_1,v_2)} + \frac{\bar{\mu}^m_{v_1,v_2}}{\sigma^{2m}_0}\right]\\ &\times \sigma^{2*m}_{v_1,v_2}.\end{aligned}$$

Given the normal prior $\mathrm{N}[\bar{\lambda}_{1,v_1,v_2}, \sigma^{2\lambda_1}_0]$, the posterior distribution of $\lambda_1(v_1, v_2)$ is

$$\begin{aligned}p(\lambda_1(v_1,v_2)|\sigma^2_\epsilon(v_1,v_2), \mu^m(v_1,v_2), \alpha(v_1,v_2), \lambda_2(v_1,v_2), \mathbf{M}_{v_1,v_2}, \mathbf{x}, \mathbf{c}_1, \mathbf{c}_2)\\ \stackrel{D}{=} \mathrm{N}[\mu^{*\lambda_1}_{v_1,v_2}, \sigma^{2*\lambda_1}_{v_1,v_2}],\end{aligned} \tag{4}$$

where

$$\begin{aligned}\sigma^{2*\lambda_1}_{v_1,v_2} &= \frac{1}{\sum_{i=1}^n c^2_{1i}/\sigma^2_\epsilon(v_1,v_2) + 1/\sigma^{2\lambda_1}_0}\\ \mu^{*\lambda_1}_{v_1,v_2} &= \left[\frac{\sum_{i=1}^n \left(m_{i,v_1,v_2} - \mu^m(v_1,v_2) - \alpha(v_1,v_2)x_i - \lambda_2(v_1,v_2)c_{2i}\right)c_{1i}}{\sigma^2_\epsilon(v_1,v_2)} + \frac{\bar{\lambda}_{1,v_1,v_2}}{\sigma^{2\lambda_1}_0}\right]\sigma^{2*\lambda_1}_{v_1,v_2}.\end{aligned}$$

Given the normal prior $\mathrm{N}[\bar{\lambda}_{2,v_1,v_2}, \sigma^{2\lambda_2}_0]$, the posterior distribution of $\lambda_2(v_1, v_2)$ is

$$\begin{aligned}p(\lambda_2(v_1,v_2)|\sigma^2_\epsilon(v_1,v_2), \mu^m(v_1,v_2), \alpha(v_1,v_2), \mathbf{M}_{v_1,v_2}, \mathbf{x}, \mathbf{c}_1, \mathbf{c}_2)\\ \stackrel{D}{=} \mathrm{N}[\mu^{*\lambda_2}_{v_1,v_2}, \sigma^{2*\lambda_2}_{v_1,v_2}],\end{aligned} \tag{5}$$

where $\sigma^{2*\lambda_2}_{v_1,v_2}$ and $\mu^{*\lambda_2}_{v_1,v_2}$ are similarly defined as those of $\lambda_1(v_1, v_2)$.

Given the inverse gamma prior $\mathrm{IG}(a^\epsilon_0, b^\epsilon_0)$, the posterior distribution of $\sigma^2_\epsilon(v_1, v_2)$ is

$$\begin{aligned}p(\sigma^2_\epsilon(v_1,v_2)|\mu^m(v_1,v_2), \alpha(v_1,v_2), \lambda_1(v_1,v_2), \lambda_2(v_1,v_2), \mathbf{M}_{v_1,v_2}, \mathbf{x}, \mathbf{c}_1, \mathbf{c}_2)\\ \sim \mathrm{IG}(a^{*\epsilon}_{v_1,v_2}, b^{*\epsilon}_{v_1,v_2}),\end{aligned} \tag{6}$$

where

$$\begin{aligned}a^{*\epsilon}_{v_1,v_2} &= a^\epsilon_0 + \frac{n}{2},\\ b^{*\epsilon}_{v_1,v_2} &= b^\epsilon_0 + \frac{\sum_{i=1}^n (m_{i,v_1,v_2} - \mu^m(v_1,v_2) - \alpha(v_1,v_2)x_i - \lambda_1(v_1,v_2)c_{1i} - \lambda_2(v_1,v_2)c_{2i})^2}{2}.\end{aligned}$$

Given the normal prior $\mathrm{N}[\gamma_0, \sigma_0^{2\gamma}]$, the posterior distribution of $\gamma$ is

$$p(\gamma|\boldsymbol{\beta}, \lambda_3, \lambda_4, \mathbf{M}, \mathbf{x}, \mathbf{c}_2, \mathbf{c}_3) \sim \mathrm{N}[\mu^{*\gamma}, \sigma^{2*\gamma}], \tag{7}$$

where

$$\sigma^{2*\gamma} = \frac{1}{\sum_{i=1}^n x_i^2/\sigma_\delta^2 + 1/\sigma_0^{2\gamma}}$$
$$\mu^{*\gamma} = \left[\frac{\sum_{i=1}^n (y_i - c_V \sum_{v_1<v_2} \beta(v_1, v_2) m_{i,v_1,v_2} - \lambda_3 c_{2i} - \lambda_4 c_{3i}) x_i}{\sigma_\delta^2} + \frac{\gamma_0}{\sigma_0^{2\gamma}}\right] \sigma^{2*\gamma}.$$

Given the normal prior $\mathrm{N}[\lambda_{30}, \sigma_{\lambda 30}^2]$, the posterior distribution of $\lambda_3$ in is

$$p(\lambda_3|\boldsymbol{\beta}, \gamma, \lambda_4, \mathbf{M}, \mathbf{x}, \mathbf{c}_2, \mathbf{c}_3) = \mathrm{N}[\mu_{\lambda_3}^*, \sigma_{\lambda_3}^{2*}], \tag{8}$$

where

$$\sigma_{\lambda_3}^{2*} = \frac{1}{\sum_{i=1}^n c_{2i}^2/\sigma_\delta^2 + 1/\sigma_{\lambda_{30}}^2}$$
$$\mu_{\lambda_3}^* = \left[\frac{\sum_{i=1}^n (y_i - c_V \sum_{v_1<v_2} \beta(v_1, v_2) m_i(v_1, v_2) - \gamma x_i - \lambda_3 c_{3i}) c_{2i}}{\sigma_\delta^2} + \frac{\lambda_{30}}{\sigma_{\lambda_{30}}^2}\right] \sigma_{\lambda_3}^{2*}.$$

Given the normal prior $\mathrm{N}[\lambda_{40}, \sigma_0^{2\lambda_4}]$, the posterior distribution of $\lambda_4$ is

$$p(\lambda_4|\boldsymbol{\beta}, \gamma, \lambda_3, \mathbf{M}, \mathbf{x}, \mathbf{c}_2, \mathbf{c}_3) = \mathrm{N}[\mu^{*\lambda_4}, \sigma^{2*\lambda_4}], \tag{9}$$

where $\mu^{*\lambda_4}$ and $\sigma^{2*\lambda_4}$ are similarly defined as those for $\lambda_3$.

### 3 Cross-validation for the tuning parameters

The tuning parameters in each of the three steps described in Section 4.3 are selected by cross-validation with $L$ folds, whereby the samples in each fold are randomly selected. Specifically:

Firstly, the tuning parameters $\{a_\alpha, b_\alpha, \sigma_{\alpha 0}^{2\theta}\}$ are chosen to minimize the quantity

$$\frac{2}{nV(V-2)(L-1)} \sum_{l=1}^L \left\{ \sum_{i \notin \text{group}_l} \left[ \sum_{v_1<v_2} (m_{i,v_1,v_2} - \hat{\mu}_l^m(v_1, v_2) - \hat{\alpha}_l(v_1, v_2) x_i \right.\right.$$
$$\left.\left. - \hat{\lambda}_{1l}(v_1, v_2) c_{1i} - \hat{\lambda}_{2l}(v_1, v_2) c_{2i})^2 \right] \right\},$$

where $\hat{\mu}_l(v_1, v_2)$, $\hat{\alpha}_l(v_1, v_2)$, $\hat{\lambda}_{1l}(v_1, v_2)$, and $\hat{\lambda}_{2l}(v_1, v_2)$ are estimated in each fold without using data from the $l$th group by fixing $c = 0$.

Secondly, tuning parameters $\{a_\beta, b_\beta, \sigma^{2\theta}_{\beta 0}, \sigma^2_\delta\}$ are chosen to minimize

$$\frac{1}{n(L-1)} \sum_{l=1}^{L} \left\{ \sum_{i \notin \text{group}_l} \left[ y_i - c_V \sum_{v_1 < v_2} \hat{\beta}_l(v_1, v_2) m_{i,v_1,v_2} - \hat{\gamma}_l x_i - \hat{\lambda}_{3l} c_{2i} - \hat{\lambda}_{4l} c_{3i} \right]^2 \right\},$$

where $\hat{\beta}_l(v_1, v_2)$, $\hat{\gamma}_l$, $\hat{\lambda}_{3l}$, and $\hat{\lambda}_{4l}$ are estimated in each fold without using data from the $l$th group while fixing $c = 0$.

Finally, fixing the rest of the tuning parameters at pre-determined values, $c$ is chosen to minimize

$$\sum_{l=1}^{L} \left\{ \sum_{i \notin \text{group}_l} \left[ \frac{2}{V(V-1)} \sum_{v_1 < v_2} (m_{i,v_1,v_2} - \hat{\mu}^m_l(v_1, v_2) - \hat{\alpha}_l(v_1, v_2) x_i - \hat{\lambda}_{1l}(v_1, v_2) c_{1i})^2 - \hat{\lambda}_{2l}(v_1, v_2) c_{2i})^2 \right. \right.$$
$$\left. \left. + (y_i - c_V \sum_{v_1 < v_2} \hat{\beta}_l(v_1, v_2) m_{i,v_1,v_2} - \hat{\gamma}_l x_i - \hat{\lambda}_{3l} c_{2i} - \hat{\lambda}_{4l} c_{3i})^2 \right] \right\} \frac{1}{n(L-1)},$$

where $\hat{\mu}_l(v_1, v_2)$, $\hat{\alpha}_l(v_1, v_2)$, $\hat{\lambda}_{1l}(v_1, v_2)$, $\hat{\lambda}_{2l}(v_1, v_2)$, $\hat{\beta}_l(v_1, v_2)$, $\hat{\gamma}_l$, $\hat{\lambda}_{3l}$, and $\hat{\lambda}_{4l}$ are estimated in each fold without using data from the $l$th group.

We end this subsection with some guidance about finding suitable ranges of tuning parameters for the cross-validation procedure. In accordance with [1], a useful starting range for the Ising prior parameters, $a_\alpha$ and $a_\beta$, is $(-4, 0)$, where $-4$ and 0 correspond sparse and dense coefficients, respectively. The suggested range for $b_\alpha$ and $b_\beta$ is $(0, 1)$, where 1 enforces the scale freeness whereas 0 enforces larger communities. We also suggest choosing $\sigma^2_\delta$ with cross-validation because it influences the variable selection of the matrix coefficients and its optimal choice depends on the scale of the connectivity mediator and the signal strength. The subnetwork smoothness of $\boldsymbol{\alpha}$ and $\boldsymbol{\beta}$ depends on $\sigma^2_{\alpha 0}$ and $\sigma^2_{\beta 0}$, respectively, the choices of which are less critical and also depend on the scale of the imaging mediator.

### 4 Additional tables

Table 1: Full names and major functions of the ROIs in main text Table 5.

| | $\hat{\boldsymbol{\alpha}}$-related | |
|---|---|---|
| ROI | Full name | Functions |
| L_FOP2 | Left Frontal Operculum 2 | Involved in language and speech processing. |
| L_STSva | Left Superior Temporal Sulcus, Ventral Anterior | Involved in social perception and audiovisual integration. |
| R_PBelt | Right Parabelt | Part of the auditory cortex, involved in processing complex sounds. |
| R_RSC | Right Retrosplenial Cortex | Involved in spatial navigation and memory. |
| L_V3 | Left Visual Area 3 | Part of the visual cortex, involved in processing visual information. |
| L_MT | Left Middle Temporal Area | Involved in processing motion and visual stimuli. |
| L_6ma | Left Supplementary Motor Area (Medial part) | Involved in planning and coordinating movement. |
| L_46 | Left Area 46 (Dorsolateral Prefrontal Cortex) | Involved in executive functions such as working memory and decision-making. |
| R_POS1 | Right Parieto-Occipital Sulcus 1 | Involved in visual and spatial processing. |
| L_8BM | Left Brodmann Area 8, Medial part | Associated with eye movements and executive functions. |

Table 2: Full names and major functions of the ROIs in main text Table 5.

| | $\hat{\boldsymbol{\beta}}$-related | |
|---|---|---|
| ROI | Full name | Functions |
| L_IP1 | Left Intraparietal Area 1 | Involved in visuospatial attention and hand-eye coordination. |
| L_VMV3 | Left Ventromedial Visual Area 3 | Involved in visual information processing related to object recognition. |
| L_TGv | Left Temporal Gyrus, ventral part | Involved in processing auditory information and language. |
| R_V3 | Right Visual Area 3 | Involved in visual processing. |
| BrStem | Brainstem | Controls vital functions and acts as a conduit for information between the brain and the body. |
| R_Thal | Right Thalamus | Acts as a relay station for sensory and motor signals to the cerebral cortex. |
| L_24dd | Left Dorsal Division of Anterior Cingulate Cortex (Area 24dd) | Involved in emotional processing and regulation. |
| L_MT | Left Middle Temporal Area (also known as V5) | Involved in processing motion and visual stimuli. |
| L_25 | Left Area 25 (Subgenual Area) | Involved in mood regulation and emotional processing. |
| L_PHT | Left Parahippocampal Gyrus, temporopolar part | Involved in memory encoding and retrieval. |

Table 3: Full names and major functions of the ROIs in main text Table 5.

| | $\hat{\boldsymbol{\alpha}} \cdot \hat{\boldsymbol{\beta}}$-related | |
|---|---|---|
| ROI | Full name | Functions |
| L_IP1 | Left Intraparietal Area 1 | Involved in visuospatial attention and hand-eye coordination. |
| L_V3CD | Left Ventromedial Visual Area 3 (CD) | Involved in high-level visual processing and object recognition. |
| L_V3 | Left Visual Area 3 | Critical for basic visual processing. |
| L_LIPv | Left Lateral Intraparietal Visual area | Important for visuospatial attention and eye movements. |
| L_a9-46v | Left Anterior Area 9-46v | Part of the prefrontal cortex, involved in working memory and executive function. |
| L_A4 | Left Auditory Area 4 | Involved in auditory processing. |
| R_TGd | Right Temporal Gyrus, dorsal part | Involved in processing auditory information and language. |
| L_47l | Left Area 47l | Involved in language and semantic processing. |
| L_25 | Left Area 25 (Subgenual Area) | Involved in mood regulation and emotional processing. |
| R_VMV2 | Right Ventromedial Visual Area 2 | Involved in visual processing, particularly in the ventral stream related to object recognition. |

Table 4: Full names of the ROIs in main text Table 6.

| | $\hat{\boldsymbol{\alpha}}$-related |
|---|---|
| Connection | Full name |
| L_VVC - L_V3A | Left Ventral Visual Cortex - Left Visual Area 3A |
| R_ProS - R_VMV1 | Right Prostriata - Right Ventromedial Visual Area 1 |
| L_V3A - L_31a | Left Visual Area 3A - Left Area 31a |
| L_IP2 - L_3b | Left Intraparietal Area 2 - Left Primary Somatosensory Cortex |
| L_PIT - L_IP1 | Left Posterior Inferior Temporal area - Left Intraparietal Area 1 |
| L_IP0 - L_31pd | Left Intraparietal Area 0 - Left Area 31pd |
| R_a9 - 46v, R_6r | Right Anterior Area 9-46v - Right Area 6r |
| L_V6 - R_V8 | Left Visual Area 6 - Right Visual Area 8 |
| R_FEF - R_RSC | Right Frontal Eye Field - RightRetrosplenial Cortex |
| L_IP2 - L_MST | Left Intraparietal Area 2 - Left Medial Superior Temporal area |

Table 5: Full names of the ROIs in main text Table 6.

| $\hat{\boldsymbol{\beta}}$-related | |
|---|---|
| Connection | Full name |
| R_Pall - R_Put | Right Pallidum - Right Putamen |
| R_FOP5 - R_7Pl | Right Frontal Operculum 5 - Right Superior Parietal Lobule |
| L_PHA3 - R_V1 | Left Parahippocampal Area 3 - Right Primary Visual Cortex |
| L_STSda - L_TPOJ2 | Left Superior Temporal Sulcus, dorsal anterior part - Left Temporo-Parieto-Occipital Junction 2 |
| R_V3 - L_PSL | Right Visual Area 3 - Left Superior Parietal Lobule |
| L_TF - R_V1 Left | Temporal Fusiform Cortex - Right Primary Visual Cortex |
| L_IP1 - L_V3CD | Left Intraparietal Area 1 - Left Ventromedial Visual Area 3CD |
| L_45 - L_Caud | Left Area 45 (part of Broca's area) and Left Caudate nucleus |
| R_V1 - L_STSda | Right Primary Visual Cortex - Left Superior Temporal Sulcus, dorsal anterior part |
| R_STGa - R_TGd | Right Superior Temporal Gyrus anterior part - Right Temporal Gyrus dorsal part |

Table 6: Full names of the ROIs in main text Table 6.

| $\hat{\boldsymbol{\alpha}} \cdot \hat{\boldsymbol{\beta}}$-related | |
|---|---|
| Connection | Full name |
| L_PIT - L_IP1 | Left Posterior Inferior Temporal area - Left Intraparietal Area 1 |
| L_V3CD - L_IP2 | Left Ventromedial Visual Area 3CD - Left Intraparietal Area 2 |
| L_PGs - L_25 | Left Parietal Gyrus superior part - Left Area 25 |
| L_IP1 - L_PEF | Left Intraparietal Area 1 - Left Frontal Eye Field |
| L_IP1 - L_VMV2 | Left Intraparietal Area 1 - Left Ventromedial Visual Area 2 |
| R_PEF - L_LIPv | Right Frontal Eye Field - Left Lateral Intraparietal Visual area |
| L_IP1 - L_V3CD | Left Intraparietal Area 1 - Left Ventromedial Visual Area 3CD |
| L_47m - L_47l | Left Area 47m - Left Area 47l |
| L_A4 - L_a9-46v | Left Auditory Area 4 - Left Anterior Area 9-46v |
| L_VMV1 - L_IP1 | Left Ventromedial Visual Area 1 - Left Intraparietal Area 1 |

### 5 Additional figures

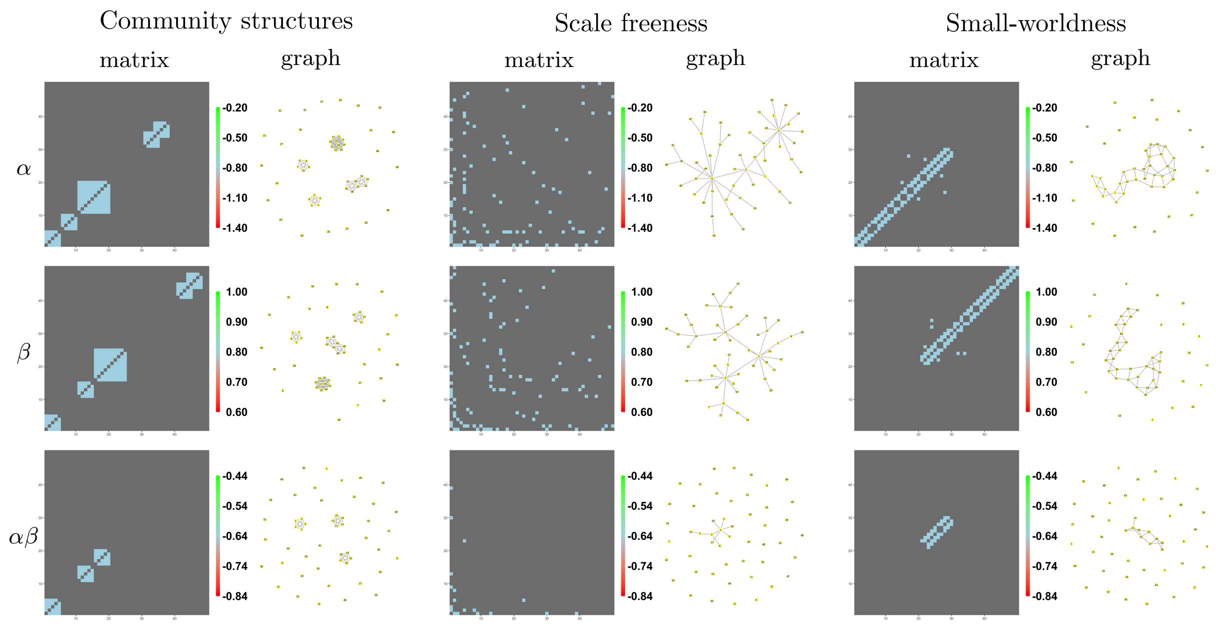


Figure 1: The network graphs corresponding to the true matrix coefficients in Section 5.1.

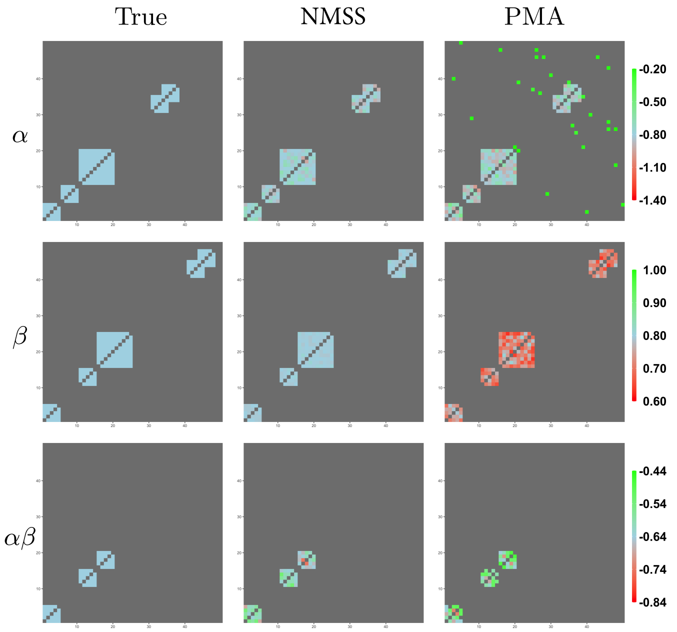


Figure 2: Heat maps of the estimated matrix coefficients produced by NMSS and PMA in one randomly picked replication in the community structures scenario.

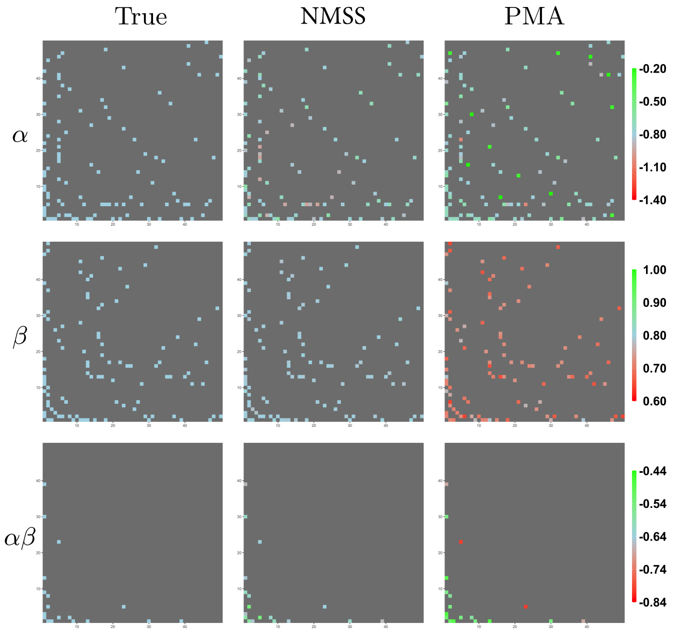


Figure 3: Heat maps of the estimated matrix coefficients produced by NMSS and PMA in one randomly picked replication in the scale freeness scenario.

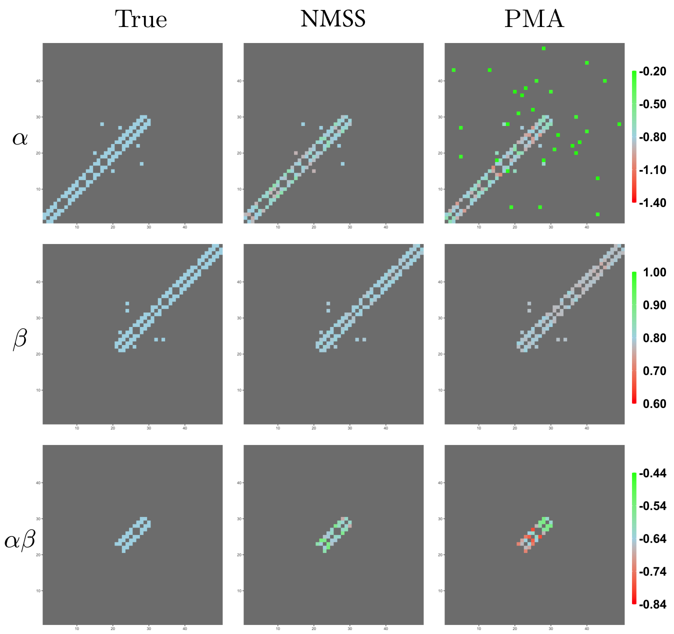


Figure 4: Heat maps of the estimated matrix coefficients produced by NMSS and PMA in one randomly picked replication in the small-worldness scenario.

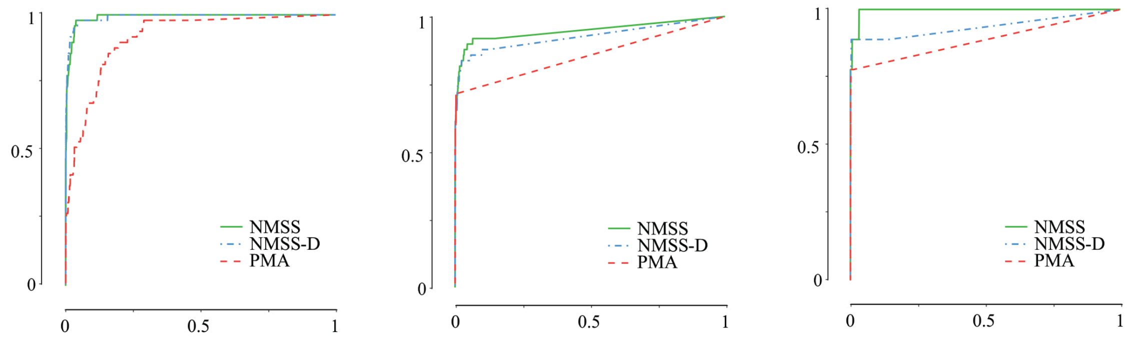


Figure 5: Comparison of ROC curves under low SNR in the scale freeness scenario across NMSS, BMSS-D, and PMA methods.

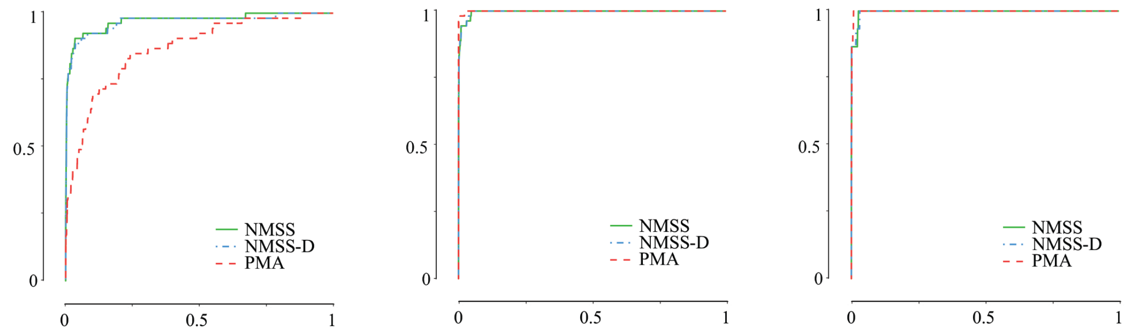


Figure 6: Comparison of ROC curves under low SNR in the small worldness scenario across NMSS, BMSS-D, and PMA methods.

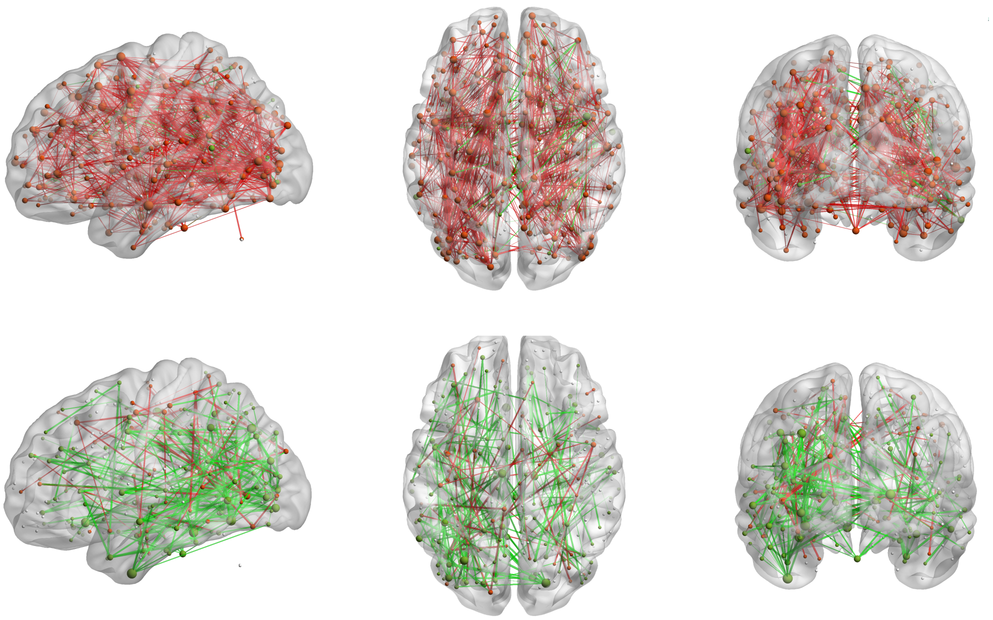

Figure 7: Visualization of the estimated matrix coefficients $\hat{\boldsymbol{\alpha}}$ (first row) and $\hat{\boldsymbol{\beta}}$ (second row) in the TOTXVI analysis.
